# Dynamically adjusted cell fate decisions and resilience to mutant invasion during steady state hematopoiesis revealed by an experimentally parameterized mathematical model

**DOI:** 10.1101/2023.12.17.572074

**Authors:** Natalia L. Komarova, Chiara Rignot, Angela G. Fleischman, Dominik Wodarz

**Affiliations:** Department of Mathematics, University of California San Diego, 9500 Gilman Drive, La Jolla, CA 92093; Department of Mathematics, University of California Irvine, Irvine CA 92697, USA; Department of Medicine, University of California Irvine, Irvine CA 92697, USA; Department of Ecology, Behavior & Evolution, University of California San Diego, 9500 Gilman Drive, La Jolla CA 92093, USA

## Abstract

A major next step in hematopoietic stem cell (HSC) biology is to obtain a thorough quantitative understanding of cellular and evolutionary dynamics involved in undisturbed hematopoiesis. Mathematical models are key in this respect, and are most powerful when parameterized experimentally and containing sufficient biological complexity. Mathematical models of hematopoiesis have either been parameterized experimentally without non-linear dynamics, or they include these complexities but have not been parameterized to the same extent. We bridge this gap using mouse data to parameterize a mathematical model of hematopoiesis that includes homeostatic control mechanisms as well as clonal evolution. We find that non-linear feedback control drastically changes the interpretation of kinetic estimates at homeostasis. This suggests that short-term HSC and multipotent progenitors (MPPs) can dynamically adjust to sustain themselves in the absence of long-term HSCs, even if they differentiate more often than they self-renew in undisturbed homeostasis. Additionally, the presence of feedback control in the model renders the system resilient against mutant invasion. Invasion barriers, however, can be overcome by a combination of age-related changes in stem cell differentiation and a mutant-associated inflammatory environment. This helps us understand the evolution of e.g. *TET2, DNMT3A*, or *JAK2* mutants, and how to potentially reduce mutant burden.

## 1. Introduction

The biology of hematopoietic stem cells has been subject to much investigation, and an increasingly detailed picture has been emerging, which includes the hierarchical structure of the hematopoietic system [1], the mechanisms underlying the associated cell fate decisions [2-6], and the kinetics of these processes [7-14]. At the same time, it has become clear that the dynamics within the hematopoietic system are characterized by complex interactions among cells, involving homeostatic control mechanisms [15,16]. Further complexity is added through evolutionary processes [17-19], resulting in the emergence of specific cell clones that can potentially increase the risk of subsequent malignant transformation and also trigger a number of non-malignant chronic health conditions [20]. Mathematical models have been used to parse the complexity of the interactions occurring within the hematopoietic system, and to investigate the principles according to which mutant cell clones emerge and give rise to tumors [21-37].

Mathematical models are most powerful when informed by experimental or clinical data. Indeed, in recent studies, mathematical models of hematopoiesis have been parameterized with the help or kinetic label progression data in mice [7], and the properties of these models have been investigated [38]. On the one hand, while these experimentally parameterized models describe the basic dynamics of cell self-renewal and differentiation, they have so far not taken into account more complex interactions that occur within the hematopoietic system, such as non-linear homeostatic control mechanisms or cellular evolutionary dynamics at homeostasis. On the other hand, mathematical models that do include these complexities [21-37] have been analyzed largely from a theoretical point of view, elucidating important principles of homeostatic control and mutant evolution. Such an analysis, however, has not been performed in the context of parameters that have been estimated experimentally, likely due to the challenges of the underlying model complexity.

Our paper addresses this gap. We calibrate a mathematical model of hematopoiesis that takes into account non-linear homeostatic control mechanisms, using existing label progression data. This approach allows us to run computer simulations using feedback control parameters that are consistent with cellular dynamics observed at homeostatic equilibrium in mice, and to interpret the meaning of experimentally estimated rates of self-renewal and differentiation among long-term hematopoietic stem cells (LT-HSCs), short-term stem cells (ST-HSCs), and multipotent progenitor cells (MPPs). We extend this model to include the dynamics of clonal evolution at homeostatic steady state and investigate the resilience of the hematopoietic system against mutant invasion, the conditions for specific mutants to emerge successfully, and which cell populations in the hierarchy of the hematopoietic system are most susceptible to mutant emergence.

## 2. Model-based interpretation of estimated self-renewal rates of ST-HSCs and MPPs

Our analysis is based on a mathematical model of hematopoiesis that has been extensively used in the literature [21-37], illustrated in Figure 1 and given by a set of ordinary differential equations that describe the dynamics of cell populations over time. We explicitly describe the dynamics of LT-HSCs (*x*_*0*_), ST-HSCs (*x*_*1*_), and MPPs (*x*_*2*_). These cells are assumed to divide with a rate *r*_*i*_. A division is assumed to result in self-renewal (two daughter stem cells) with a probability *p*_*i*_, and in differentiation (two daughter downstream cells) with a probability *1-p*_*i*_. We also include populations of common myeloid progenitor (CMP, *x*_*3*_^*(1)*^) and common lymphoid progenitor (CLP, *x*_*3*_^*(2)*^) cells.

**Figure 1:**
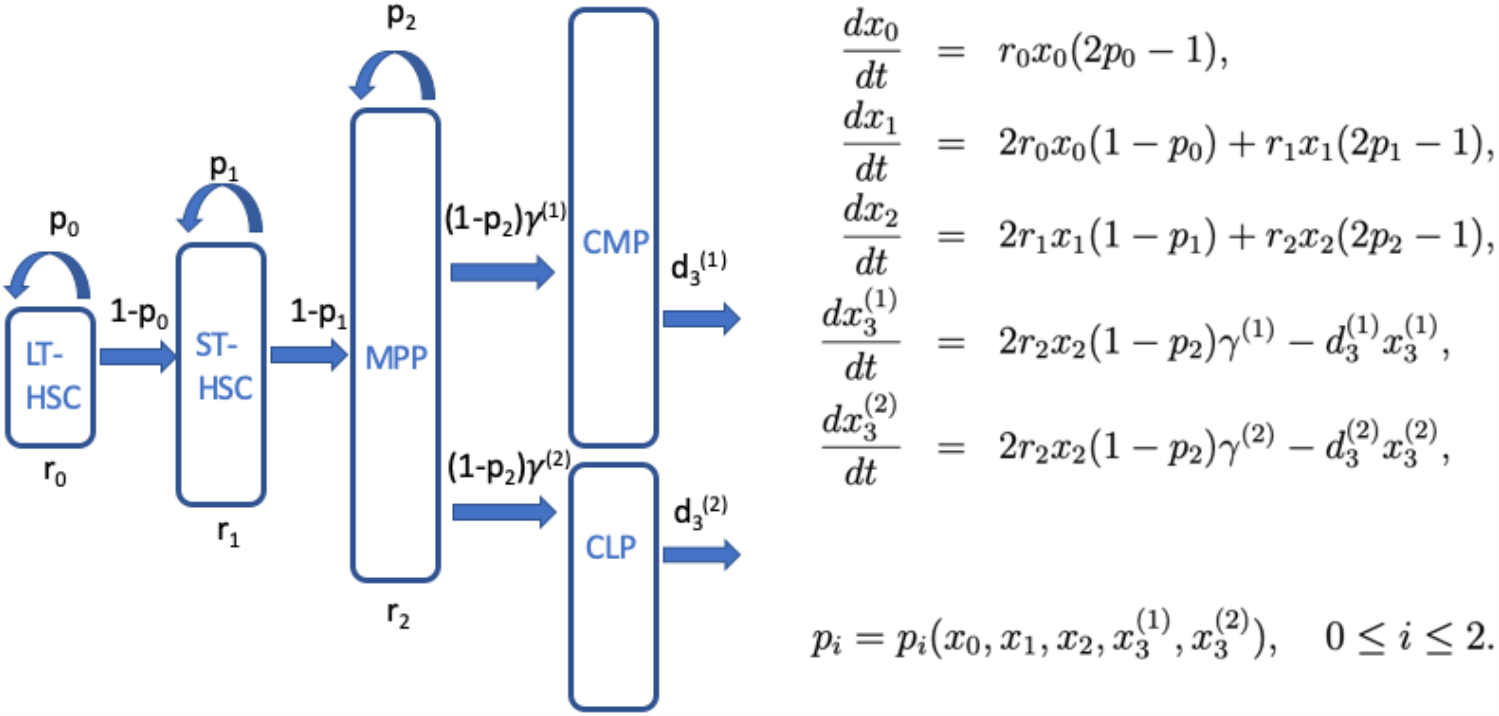
A schematic of the mathematical model showing the four compartments and the associated division rates and self-renewal probabilities. The system of ordinary differential equations used to describe the dynamics is provided on the right.

This model is parameterized with the help of existing neutral label propagation data in mice at homeostatic setpoint, first published by Busch et al [7], and expanded with additional data in a subsequent study [39]. There, 1% of LT-HSCs was labeled, and the progression of the neutral label in ST-HSCs, MPPs, CMPs, and CLPs was documented for about 450 days. The model was fit to these data as described in the Supplementary Materials, and the best fitting parameters as well as their confidence intervals are given in Table 1. The fits and the data are presented in Supplementary Figure S2. A similar model fitting approach was taken to estimate parameters by Busch et al [7], but the model we use is structurally different (due to our focus on evolutionary dynamics), thus necessitating the parameter estimation procedures performed here.

**Table 1:** Definitions of model parameters and their values. The bar over values 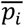 and 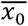 is used to indicate that these are constants, and to distinguish these from functions *p*_*i*_*(x*_*0*_,*…)* and *x*_*0*_*(t)*. * There is a degree of uncertainty about the number of LT-HSCs (7), but none of the qualitative results depend on this value: all the values simply scale with the number of LT-HSCs and a change in this value would result in a multiplicative factor.

| Notation | Parameter definition | Value | 95% C.I. | Units |
| --- | --- | --- | --- | --- |
| $r_0$ | Division rate of LT-HSCs | 0.0107 | (0.0084, 0.014) | days<sup>-1</sup> |
| $r_1$ | Division rate of ST-HSCs | 0.067 | (0.021, 0.10) | days<sup>-1</sup> |
| $r_2$ | Division rate of MPPs | 0.136 | (0.066, 0.95) | days<sup>-1</sup> |
| $\bar{p}_0$ | Self-renewal probability of LT-HSCs | 0.5 | | 1 |
| $\bar{p}_1$ | Self-renewal probability of ST-HSCs | 0.473 | (0.40, 0.48) | 1 |
| $\bar{p}_2$ | Self-renewal probability of MPPs | 0.416 | (0.323, 0.49) | 1 |
| $d_3^{(1)}$ | Removal rate of CMPs | 0.034 | (0.017, 0.22) | days<sup>-1</sup> |
| $d_3^{(2)}$ | Removal rate of CLPs | 0.007 | (0.005, 0.017) | days<sup>-1</sup> |
| $\bar{x}_0$ | Equilibrium number of LT-HSCs | 17,000* | | cells |

We will first briefly consider the scenario where the estimated parameters are constants. We subsequently examine the dynamics under the assumption that the estimated parameters (in particular, the cells’ probability of self-renewal) at homeostasis are subject to feedback control.

### 2.1 Constant parameter values

In the most basic setting, we can assume that the rate of cell division and self-renewal at homeostasis are constants, “programmed” into the cells. Hence, we can use the estimated parameter values to simply set 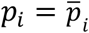 (Table 1). Because the self-renewal probability of LT-HSCs is assumed to be exactly one half 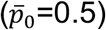, the model is characterized by a stable equilibrium that can describe homeostasis (see Supplement, Section 1). The estimated self-renewal rates of the downstream compartments are both below one half 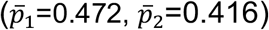. Under the current assumptions, this means that the ST-HSC and the MPP compartments cannot maintain themselves, and that their persistence strictly depends on the input from the LT-HSCs. This has also been the conclusion of [7].

### 2.2. Parameter values are determined by feedback control at homeostasis

An alternative, and biologically more complex assumption is that the observed equilibrium rates of cell division and self-renewal are maintained dynamically through feedback control mechanisms. Experimental data indicate that both the rate of cell division, *r*_*i*_, and the probability of self-renewal, *p*_*i*_, might be subject to feedback control, because these processes seem to accelerate during tissue reconstitution compared to homeostatic setpoint [7]. Here we assume that the self-renewal probability, *p*_*i*_, of LT-HSCs, ST-HSCs, and MPPs can be influenced by feedback regulation. For simplicity, we continue to treat the division rate of cells, *r*_*i*_, as constants. The reason is that in the current study we focus on dynamics at homeostasis, and in this context, only assumptions about the self-renewal probability of cells have an impact on the stability properties of the model [40].

#### Modeling feedback control

Since the exact functional form of feedback on the self-renewal probability, *p*_*i*_, is not known, different possibilities have been explored in the literature. For example, in previous studies [21-23] it was assumed that the final, most differentiated cell population reduces the self-renewal probability of cells (importantly *p*_*0*_), which then results in the presence of stable equilibria. There are many other possible ways in which the self-renewal probability of cells could be regulated. We could write in general that in compartment *i*, the probability of self-renewal of cells is some function of cell populations in the same and other compartments: *p*_*i*_*=p*_*i*_*(x*_*0*_,*x*_*1*_,*x*_*2*_,*x*_*3*_). At healthy, homeostatic equilibrium, the value of this function is assumed to be equal to the experimentally measured self-renewal probability, 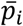 (Table 1).

Here, we explore how the interpretation of the estimated self-renewal probabilities can change if we assume that homeostasis is driven by feedback control. In the Supplement, we provide calculations using different types of control. We show that results reported here hold true generally, for a large class of feedback regulation functions that satisfy some mild conditions (such as the existence of a stable equilibrium). Here, for illustration purposes, we will discuss results in the context of one specific feedback loop, where homeostasis is maintained by feedback of LT-HSCs on their own self-renewal probability, *p*_*0*_, and similar feedback by populations of the downstream compartments on their own self-renewal probability. Mathematically, this is expressed by assuming that *p*_*0*_ is a decreasing function of *x*_*0*_, for example, *p*_*0*_*=c*_*0*_*/(h*_*0*_*x*_*0*_*+1)*, where *c*_*0*_ is the LT-HSC self-renewal probability in the absence of feedback, and *h*_*0*_ is the strength of the feedback regulation. While this alone is sufficient to achieve a stable equilibrium, we will assume that the ST-HSC and MPP cell populations (*x*_*1*_ and *x*_*2*_) can potentially limit their own self-renewal probabilities in a similar way, i.e. *p*_*1*_*=c*_*1*_*/(h*_*1*_*x*_*1*_*+1)* and *p*_*2*_*=c*_*2*_*/(h*_*2*_*x*_*2*_*+1)*. While this formulation can correspond to feedback mediated by the cells themselves, it can also be interpreted to represent feedback from the microenvironment, which senses the number of cells.

#### Model calibration in the presence of feedback

Using our estimated parameters (Table 1), we have *p*_*0*_*=c*_*0*_*/(h*_*0*_*X*_*0*_*+1)=0*.*5*, where *X*_*0*_ is the LT-HSC population at homeostasis. It has been estimated that a mouse contains approximately 1.7×10^4^ hematopoietic stem cells [7] (our conclusions do not depend on the accuracy of this number, see Table 1 and Supplementary Materials). For an assumed (and unknown) value of *h*_*0*_, which quantifies the amount of regulation that takes place in the system (the higher *h*_*0*_, the stronger the control), we determine the value of *c*_*0*_ such that the LT-HSC population equals 1.7×10^4^. Similarly, setting the values of *h*_*1*_ and *h*_*2*_ to a desired level (describing the strength of the control loops for ST-HSCs and MPPs), and using the equations 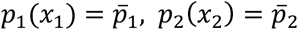, we can calculate the values of *c*_*1*_ and *c*_*2*_ such that the effective self-renewal probabilities at equilibrium matches the experimentally observed ones. The following choices guarantee that at the equilibrium, the self-renewal probabilities coincide with their experimentally measured values for LT-HSCs, ST-HSCs, and MPPs:

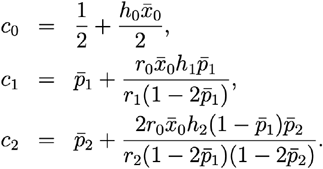

(For calibration under more general assumptions on the feedback function, see Supplement Section 1). Note that if we set *h*_*i*_*=0* (no control, constant self-renewal probabilities), we simply obtain that the self-renewal probabilities *p*_*0*_, *p*_*1*_, and *p*_*2*_ are equal to the experimentally measured values 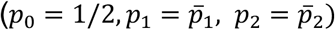.

#### Can ST-HSCs and MPPs self-maintain?

Given that the estimated self-renewal probabilities of ST-HSCs and MPPs are less than one-half (Table 1), the first interpretation that comes to mind is that these populations cannot maintain themselves in the absence of input from the LT-HSC compartment. According to the model with feedback control, however, this need not be the case. Let us consider ST-HSCs as an example. Let us assume that the self-renewal probability of ST-HSC in the absence of negative feedback inhibition, *c*_*1*_, is greater than one half. This means that the ST-HSC cell population can maintain itself even without the input from LT-HSCs. At the same time, in the context of our model, the negative feedback will result in an equilibrium value of p_1_<0.5 (and equal to the experimentally measured rate of self-renewal, 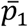), giving the false impression that this cell compartment cannot persist in the absence of LT-HSCs.

A biological interpretation of the above mathematical statement is as follows. The experimentally measured effective self-renewal probability of ST-HSCs and MPPs (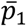 and 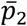) at equilibrium is determined by both (i) the basic division dynamics of the cells in these compartments and (ii) the influx of cells from the upstream compartment. If there is no influx from cells upstream in the differentiation pathway (as is the case for LT-HSCs), the self-renewal probability at equilibrium in the presence of negative feedback is exactly one half, that is, at the population level, half of the cell divisions result in self renewal, and half result in downstream differentiation. The presence of cell influx from an upstream compartment, however, changes this balance, reducing the effective self-renewal probability at equilibrium to a value less than one half. The stronger the cell influx from the upstream compartment, the lower the equilibrium self-renewal probability falls below one half. This is because the influx increases the number of cells in this compartment beyond what division events alone would achieve, thus raising the overall amount of negative feedback.

Since cells now differentiate more often than self-renew, it gives the impression that these cell populations cannot maintain themselves. It follows from our model, however, that if the upstream compartment is removed and the cell influx stops, the cell population that previously differentiated more often than it self-renewed might now effectively act like a stem cell compartment with equal self-renewal and differentiation rates. This is shown in Figure 2. The simulation starts at homeostasis, where population sizes are at equilibrium and the self-renewal probability of LT-HSCs is one half, while the self-renewal probability of ST-HSCs and MPPs is less than one half, using the estimated parameters (Table 1). At the indicated time point, the LT-HSCs (blue line) are removed in Figure 2A. Consequently, the cell populations adjust but do not go extinct (Figure 2A, top). The self-renewal probability of the ST-HSCs rises to exactly one half (Figure 2A, bottom), because the influx of LT-HSCs due to differentiation has stopped. The self-renewal probability of MPP also increases, but remains below one half, due to the continued influx of ST-HSCs. In Figure 2B, the same kind of simulation is repeated, but both LT-HSCs and ST-HSCs are depleted, as indicated. Now, the self-renewal probability of MPPs rises to exactly one half, due to the lack of influx from upstream cell populations, allowing the MPPs to maintain the system.

**Figure 2.**
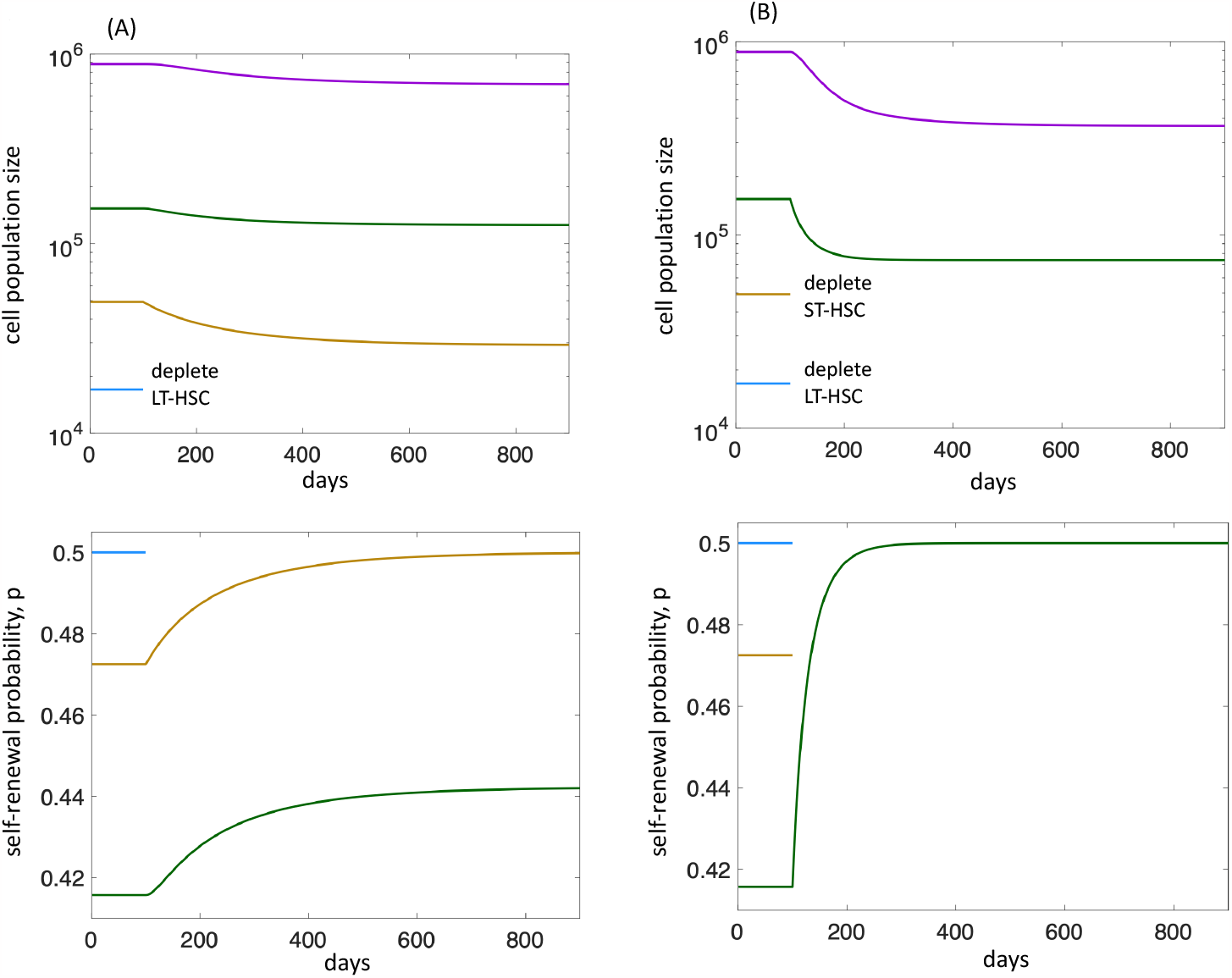
Computer simulation of stem cell depletion. LT-HSCs, ST-HSCs, MPPs, and the sum of CMPs and CLPs are shown in blue, yellow, green, and purple lines. Initially, the system is at homeostatic equilibrium. (A) At 100 days, the LT-HSC population is removed. (B) At 100 days, both the LT-HSCs and the ST-HSCs are removed. We note that this simulation does not include replication limits. If replication limits were included, population extinction would eventually occur when the cells have exhausted their replicative capacity. Negative feedback on the self-renewal probabilities by cells in the same compartment was used, with *h*_*0*_=10^−5^, *h*_*1*_=10^-5.5^, *h*_*2*_=10^-5.5^. The rest of the parameters are given in Table1.

The same result can be derived for a more general functional form of feedback control (Supplementary Materials). The important point is that in models with control that are consistent with experimentally measured parameters, compartments that exhibit a self-renewal probability <1/2 may still be able to self-maintain, due to the adjusted increase of this probability in the absence of an upstream compartment.

We note that the ability of these downstream cell populations to “maintain themselves” indefinitely in the absence of LT-HSCs in our model is due to the absence of replication limits in the equations that describe the dynamics of these cell populations. If, however, down-stream cell populations are subject to replication limits (see Supplement, Section 4), maintenance of these cell populations in the absence of LT-HSCs is only temporary. Once their replicative potential is exhausted, the cell populations are predicted to go extinct. These model properties are consistent with experimental data in which ST-HSCs or MPPs can at least temporarily maintain a functional hematopoietic system even if the LT-HSC population is compromised [41-44].

## 3. Evolutionary Dynamics in the estimated parameter space

Next, we investigate the evolutionary dynamics in this model using the estimated parameters. Denoting wild-type LT-HSCs, ST-HSCs, MPPs and further downstream CMP and CLP cells by *x*_*0*_, *x*_*1*_, *x*_*2*_, *x*_*3*_^*(1)*^, and *x*_*3*_^*(2)*^, respectively, and the corresponding mutant cells by *y*_*0*_, *y*_*1*_, *y*_*2*_, *y*_*3*_^*(1)*^ and *y*_*3*_^*(2)*^, the model with feedback control is given by the following set of ODEs:

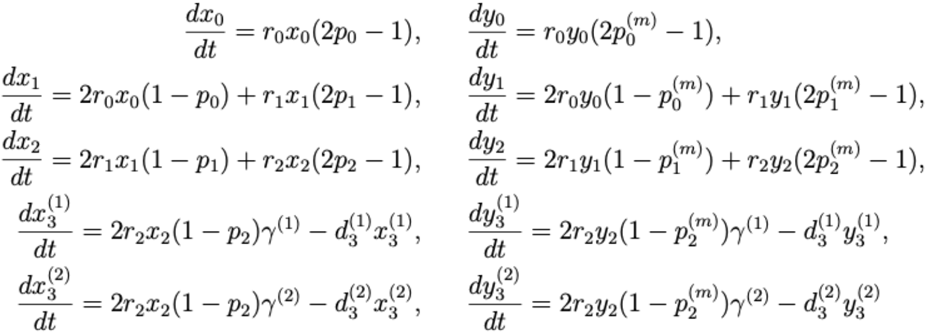

Mutant-specific parameters are denoted by superscript (m). WT and mutant cells are assumed to compete with each other through shared feedback control, e.g. the self-renewal probability for WT and mutant LT-HSCs is given by *p*_*0*_*=c*_*0*_*/(h*_*0*_*(x*_*0*_*+y*_*0*_*)+1)* and p_0_^(m)^= *c*_*0*_^*(m)*^*/(h*_*0*_^*(m)*^*(x*_*0*_*+y*_*0*_*)+1)*, respectively, and similarly for ST-HSCs and MPPs. We will study the invasion dynamics of an advantageous mutant assuming that it arises in different cell compartments, including LT-HSCs, ST-HSCs, and MPPs. In the above system, de-novo mutations are not included (see Supplement, section 2 for equations describing continuous mutant generation). Instead, here we investigate the fate of mutants by placing them in different compartments and studying the resulting competition dynamics between wild-type and mutant cells, asking under which conditions the mutant can invade from low numbers.

We focus on advantageous mutants. In general, mutants may have properties different from those of wild-type cells, including an increased replication rate, an increased probability of self-renewal, or an increased replication limit. It has been shown [22,37,40] that in models of the type employed here, a mutant whose replication rate is increased (but the probability of self-renewal is unaffected) does not behave as an advantageous mutant. Therefore, in what follows we will assume that mutants’ overall replication rate is the same as that of the wild type, but that parameters determining the self-renewal probability differ.

### 3.1 Invasion barriers for mutants originating in different compartments

#### LT-HSC compartment

If the mutant emerges in the LT-HSC compartment, the evolutionary dynamics have been investigated previously in a similar model [22,37]. Assume that mutants are characterized by an increased probability of self-renewal. Suppose that the basic probability of self-renewal in the absence of feedback, *c*_*0*_, is the only parameter that varies between wild-type and mutant cells, and denote the equivalent mutant parameter by *c*_*0*_^*(m)*^. Then the condition for mutant invasion is *c*_*0*_^*(m)*^ *> c*_*0*_. The cell population with the larger basic self-renewal probability wins and replaces the competing cell population. Another parameter that determines fitness in our model is the strength of feedback inhibition of self-renewal, denoted by *h*_*0*_ in the LT-HSC compartment. If this is the only parameter that varies between WT and mutant cells, then a mutant invades and replaces the wild-type population if *h*_0_^*(m)*^*<h*. In the most general case, where the mutant self-renewal probability is given by 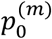, the condition for the mutant winning is given by 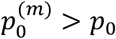, where both quantities are measured at the wild-type equilibrium. This straightforward result is consistent with the previous literature [22].

#### Downstream replicating compartments

If the mutant arises in either the ST-HSC or the MPP compartments, the conditions for mutant invasion are different. Although mutant and WT cells compete with each other in these compartments, the mutant does not necessarily invade if it has a reproductive fitness advantage (e.g., a higher self-renewal probability compared to the wild type). Instead, the reproductive advantage of the mutant needs to be sufficiently large and lie above a threshold for invasion to be successful. Let us suppose that the self-renewal rate of mutants in the ST-HSC compartment differs from that of wild-types by a constant factor: 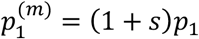, where *s* is the selection coefficient. The condition for the mutant to succeed (that is, to establish a nonzero presence) in the ST-HSC compartment is given by: 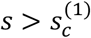, where 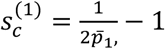 (note that this condition is general and does not depend on the specifics of the feedback function, see Supplement Section 2).

For the parameter values in Table 1, the invasion threshold for the ST-HSC compartment is approximately 0.0582. This means that if a mutant is generated in the ST-HSC compartment, its fitness advantage must exceed 5.82% for those cells to be able to establish a presence among the ST-HSC population, and subsequently invade the downstream compartments. Similarly, if a mutant originates in the MPP compartment, its fitness threshold under parameter values in Table 1 is given by 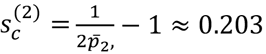, that is, fitness advantage must exceed 20.3%. At the same time, note that confidence intervals for the estimates can be relatively large (especially for MPPs), indicating a degree of uncertainty regarding the exact magnitude of the invasion threshold.

The reason for the threshold behavior is as follows. For the sake of the argument, assume that mutant cells originate in the ST-HSC compartment. In this case, they have an inherent disadvantage compared to the wild-type ST-HSCs, despite their larger self-renewal potential. The fitness of the wild-type cells in the ST-HSC compartment is determined by the combination of cell influx from the LT-HSCs (through differentiation) and the self-renewal of the wild-type ST-HSCs. In contrast, the fitness of the mutant ST-HSCs is determined only by their self-renewal rate (because no mutants are present among the upstream LT-HSCs). To overcome this inherent disadvantage of the mutant, the self-renewal potential of the mutant must be sufficiently large relative to that of the wild-type among the ST-HSCs for the mutant cells to invade there.

Figure 3 illustrates these concepts with computer simulations (see Supplement Section 3 for details of the mathematical analysis). The simulations start with the wild-type cell population at equilibrium (solid lines), and a small amount of advantageous mutant cells (dashed lines) are introduced. First, a single mutant characterized by a 2% advantage is introduced among the LT-HSCs (Figure 3A). Over time, the mutants invade, although this takes a time frame longer than the life-span of a laboratory mouse (2.5 years) under the estimated parameters. Notably, although the mutant was introduced among LT-HSCs, it first rises among the downstream cells, because differentiation propagates the mutant into these compartments, where cell reproduction occurs with a faster rate. In Figure 3B, the same kind of mutant (2% advantage) is placed among the MPP cells. Due to the invasion barrier, however, the mutants fail to invade. In contrast, if a mutant with a very large advantage is placed among the MPPs (e.g. 25% advantage, Figure 3C), the invasion barrier is overcome and the mutant invades within a relatively short time span.

**Figure 3:**
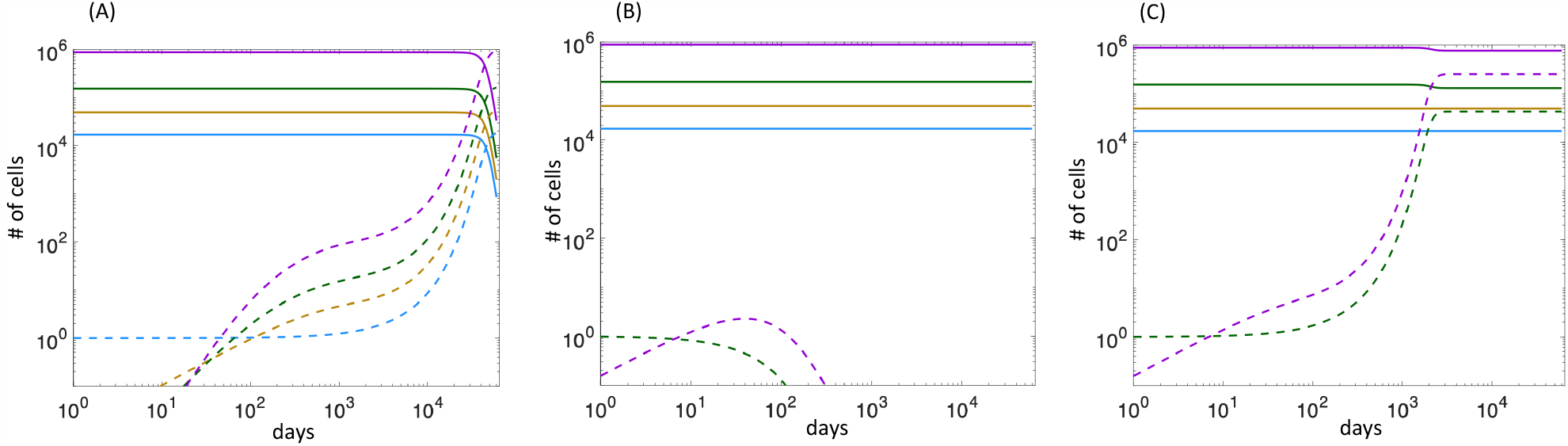
Simulated invasion dynamics of mutant cells. LT-HSCs, ST-HSCs, MPPs, and the sum of CMPs and CLPs are shown in blue, yellow, green, and purple lines, respectively. Solid line are wild-type cells, dashed line mutants. (A) A mutant with a 2% advantage is placed among LT-HSCs. (B) A mutant with a 2% advantage is placed among MPPs. (C) A mutant with a 25% advantage is placed among MPPs. Negative feedback on the self-renewal probabilities by cells in the same compartment was used, with *h*_*0*_=10^−5^, *h*_*1*_=10^-5.5^, *h*_*2*_=10^-5.5^. The rest of the parameters are given in Table1.

### 3.2 Evolutionary mechanisms to break the mutant invasion barrier in downstream compartments

Let us consider mutant invasion in the ST-HSC cell population. According to the best fitting parameter estimate, a mutant would have to enjoy a >5.8% advantage to invade from low numbers. While possible for certain types of mutants, smaller fitness advantages (e.g. 2% or less) are more common. Here we explore mechanisms that could lead to the emergence of mutants with a relatively small fitness advantage.

According to our model, the mutant invasion barrier arises due to the influx of wild-type cells from upstream compartments through differentiation. A mutant that arises downstream, e.g. among ST-HSCs, lacks that influx, conferring an inherent disadvantage. Therefore, one way to reduce the mutant invasion barrier is to reduce the influx of wild-type cells from upstream compartments. This is what might occur during the aging process. Experimental data indicate that aging in mice results in a reduced rate at which HSCs divide and differentiate [8,45,46]. In terms of our model, this can be expressed by an age (time)-dependent reduction of *r*_*0*_*x*_*0*_. The model then predicts that an advantageous mutant arising among ST-HSCs or MPPs, which might not be able to invade at a younger age, could successfully invade at an older age when the influx of upstream wild-type cells is diminished.

This is illustrated by computer simulations in Figure 4, using our base model where cells in each compartment exert negative feedback on their own self-renewal probabilities. A mutant with a 2% advantage is placed into the ST-HSC population at equilibrium. In a basic simulation, the mutant fails to grow due to the calculated invasion barrier (Figure 4A). In Figure 4B, the simulation is run such that the influx of wild-type cells from the LT-HSC compartment through differentiation (proportional to *r*_*0*_*x*_*0*_) is reduced by a small amount every time unit. Once the influx rate *r*_*0*_*x*_*0*_ falls below a threshold, the mutant cells start to grow from low numbers, meaning that the invasion barrier has been sufficiently reduced. At the same time, however, the mutant grows slowly and does not reach significant levels within the life-span of a laboratory mouse (about 2.5 years) in this parameter regime.

**Figure 4.**
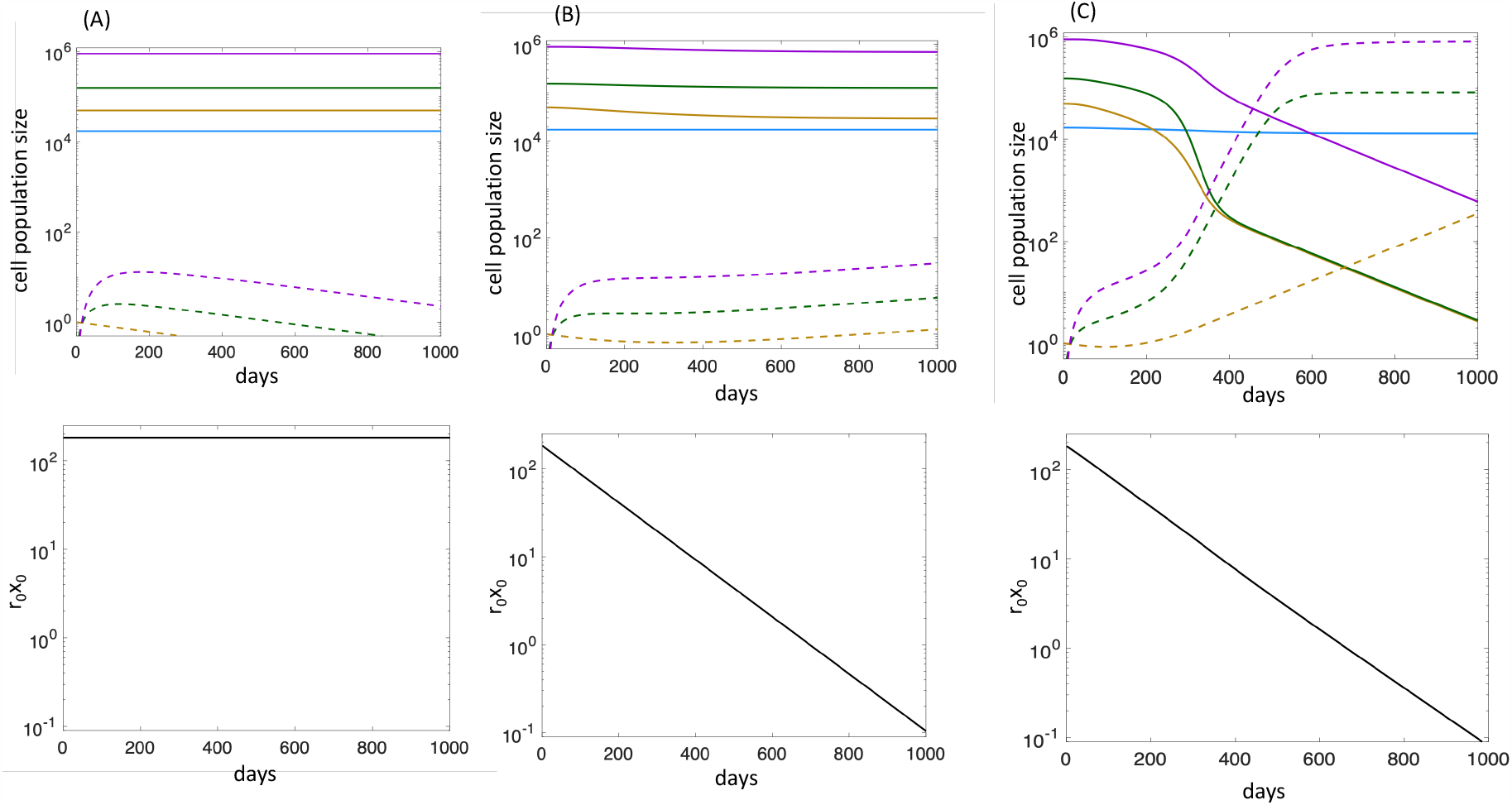
Computer simulations showing how mutants might overcome the invasion barrier in our model. A small amount of mutant cells is placed among ST-HSCs, while the wild-type cells are at equilibrium. In the top graphs, the dynamics of cell populations are shown, where LT-HSCs, ST-HSCs, MPPs, and the sum of CMPs and CLPs are shown in blue, yellow, green, and purple lines. Wild-type cells are shown by solid lines, mutant cells by dashed lines. The bottom graphs show the influx rate of LT-HSCs to ST-HSCs through differentiation (r_0_x_0_) over time. (A) model simulation without added mechanisms. The mutants fail to invade due to the barrier. (B) Aging-induced reduction of the rate of wild-type influx from LT-HSCs to ST-HSCs is modeled, by reducing the value of *r*_*0*_*x*_*0*_ by a factor of 1.0075 every day. Once the value of *r*_*0*_*x*_*0*_ has been sufficiently reduced, the mutant can invade, albeit slowly. (C) In addition to the reduction of *r*_*0*_*x*_*0*_, we also assume that the most differentiated mutant cells in our model, *y*_*3*_^*(1)*^ *+y*_*3*_^*(2)*^, induce inflammation, which reduces the self-renewal probability of wild-type cells in all compartments, but not of mutant cells. Hence, *p*_*0*_*=c*_*0*_*/(h*_*0*_*(x*_*0*_*+y*_*0*_*)+η(y*_*31*_*+ y*_*32*_*)+1), p*_*1*_*=c*_*1*_*/(h*_*1*_*(x*_*1*_*+y*_*1*_*)+η(y*_*3*_^*(1)*^*+ y*_*3*_ ^*(2)*^*)+1), p*_*2*_*=c*_*2*_*/(h*_*2*_*(x*_*2*_*+y*_*2*_*))+η(y*_*3*_^*(1)*^*+ y*_*3*_^*(2)*^*)+1)*, where *η* denotes rate at which wild-type cells are inhibited by mutant-induced inflammation. Now, mutant cells can emerge on a relatively fast time scale. We assumed that *η=0*.*01*. Negative feedback on the self-renewal probabilities by cells in the same compartment was used, with *h*_*0*_=10^−5^, *h*_*1*_=10^-5.5^, *h*_*2*_=10^-5.5^. The rest of the parameters are given in Table1.

Another mechanism that might contribute to overcoming the mutant invasion barrier is based on the observation that certain mutants in the hematopoietic system (such as *JAK2, TET2*, or *DNMT3A* mutants) are associated with an increased inflammatory environment [47,48]. Data indicate that inflammation might negatively influence the self-renewal capacity of wild-type cells, while affecting mutant cells to a lesser extent or not at all [49,50]. Indeed, it is thought that differentiated mutant cells themselves can induce increased inflammation [51,52], thus turning conditions in favor of the mutant lineage (evolutionary niche construction). We incorporated this into our mathematical model by assuming that the self-renewal probability of wild-type (but not mutant) LT-HSCs, ST-HSCs, and MPPs is reduced proportionally to the number of mutant cells in the most differentiated compartments in our model (see Figure 4 for details of the formulation). We simultaneously assumed that the influx of wild-type cells from LT-HSCs to ST-HSCs is reduced due to aging, as described above. In this simulation, the reduction of wild-type influx from LT-HSCs to ST-HSCs eventually allows the mutant cell population to grow. First, this growth is slow as before (Figure 4C). Over time, however, mutant cell growth accelerates due to a positive feedback effect. An increase in the mutant population leads to increased inflammation levels, which in turn raises the relative fitness of mutants, and hence accelerates their growth. Through a combination of the reduced wild-type cell differentiation from LT-HSCs to ST-HSCs and mutant-induced inflammation, we can observe a significant rise in mutant abundance during the life-span of a laboratory mouse (Figure 4C). Note that although the mutant was placed among the ST-HSCs, mutant invasion is predicted to be visible first in the downstream cell populations, due to their higher reproduction rates.

If the extent of mutant-induced inflammation is very strong, such that one mutant cell can already significantly increase the amount of inflammation, then the reduction of wild-type cell influx from LT-HSCs to ST-HSCs might not be needed to observe successful mutant growth. This, however, is probably not biologically realistic, because a certain amount of mutant growth is likely needed for them to sufficiently amplify inflammation levels. A generally high background level of chronic inflammation (e.g. due to an underlying disease or condition), however, might enable successful mutant invasion in the absence of an age-related reduction in wild-type cell influx from LT-HSCs to ST-HSCs.

We point out that longer-term sustained mutant presence might further require an increase in the cells’ replicative capacity. As shown in Supplementary Materials section 4, mutant invasion can be temporary if a mutant cell arising downstream (e.g. among ST-HSCs or MPPs) has limited replication capacity. In this case, further mutations might be needed to extend the replication capacity of mutant cells, thus enabling their long-term persistence. There is evidence that *tet2* and *dnmt3a* mutants already have an extended replicative capacity in mice [53,54], meaning that with these mutants, no further increase in the cellular replication capacity might be needed for long-term mutant success.

## 4. Discussion and Conclusion

Quantitative data about the kinetics of cell division, self-renewal, and differentiation in LT-HSCs, ST-HSCs, and MPPs in mice (e.g. arising from label propagation experiments [7]) represent an invaluable source for the parameterization of mathematical models that describe hematopoiesis. Our analysis, however, has shown that the interpretation of these estimated parameters can change drastically if the mathematical models include non-linear homeostatic feedback control mechanisms. Without explicitly implementing homeostatic control mechanisms in mathematical models, the parameter estimates appear to suggest that while the LT-HSC population can maintain itself because self-renewal and differentiation processes are exactly balanced at homeostasis, this is not the case for ST-HSCs or MPPs: among those cells, differentiating divisions occur more frequently than self-renewal divisions at equilibrium, implying that in the absence of LT-HSCs, these downstream populations will wash out. If feedback control is assumed to operate in the ST-HSC and MPP compartments, however, the dominance of differentiation divisions is compatible with the ability of these downstream cells to maintain themselves during homeostasis, even in the absence of LT-HSCs. The reason is that in the model, the dominance of differentiation divisions is the consequence of increased negative feedback on self-renewal that results from the influx of upstream cells (e.g. LT-HSC) through differentiation. If the upstream cells are removed, this feedback will be weaker, leading to an exact balance between self-renewal and differentiation events. The model can thus reconcile the observation that transplanted ST-HSCs and MPPs (in the absence of upstream cells) can reconstitute a partial hematopoietic system and maintain it at homeostasis, before these cells reach their replication capacity [41-43]. Similarly, following HSC and progenitor ablation in mice, HSC did not recover beyond 10% of normal numbers, while the progenitor cell population rebounded quickly and steady state hematopoiesis was for the most part not disrupted [44].

The model further suggests that feedback regulation in the ST-HSC and MPP compartments has important implications for the potential of advantageous mutants to emerge there. These compartments are thought to be important for mutant evolution because the cells replicate more frequently than LT-HSCs. Hence, chances to generate mutants are higher among ST-HSCs and MPPs compared to LT-HSCs. Due to the influx of wild-type cells from upstream compartments, however, competition among ST-HSCs and MPPs is more challenging for mutants. If a mutant is generated in one of those populations, the mutant competes with wild-type cells on two levels: (i) Through cell replication / loss dynamics within the compartment. If the mutant has a higher self-renewal rate than the wild-type, it will be advantageous in this respect. (ii) Through cell influx from the upstream compartments through differentiation. The mutant will be disadvantageous in this respect if it arose among the ST-HSCs or MPPs, because WT cells experience an influx from upstream compartments, while this is not the case for mutants (since they were only generated downstream). Therefore, to be able to spread in this setting, the replicative advantage of the mutants must be high enough to overcome this mutant disadvantage. We have derived a simple formula for the mutant invasion threshold, in which the selection coefficient, s, of mutants must be higher than *1/(2p)-1*, where *p* is the equilibrium self-renewal probability of the WT cells in the mutant’s compartment of origin.

In contrast, mutants that arise among the LT-HSCs can grow to dominance in this model if they have any degree of advantage compared to the wild-type cells, no matter how small (note that setting p=1/2 in the formula above gives a zero threshold). The reason is that there are no upstream compartments, and hence no influx of wild-type cells. However, LT-HSCs are less likely to produce mutants due to infrequent cell divisions. Therefore, the model suggests that there is a tradeoff: It is unlikely for mutants to be created among LT-HSCs, but if they do emerge they can invade if they are advantageous. In contrast, it is more likely for mutants to be generated downstream, but the extent of the fitness advantage must lie above a threshold for them to be able to take-off.

With respect to the mutant invasion threshold calculated for the best fitting parameters (Table 1), it is important to keep in mind the confidence intervals. In particular, the estimate for the self-renewal probability of MPPs is characterized by relatively large confidence intervals, meaning that there is a degree of uncertainty regarding the exact magnitude of this invasion threshold (which was estimated to be around 20% from the best-fitting parameters). Hence, it is possible that this invasion threshold might be lower, and therefore more likely to be overcome by mutated cells. Even if a mutated cell does overcome the invasion threshold, however, our simulations indicate that mutant invasion would occur on a rather slow time scale relative to the life-span of the organism. According to the parameterized model, robust mutant invasion within a biologically meaningful time frame would require the mutant advantage to be significantly larger than the invasion threshold (even for mutants emerging among MPPs), again pointing towards a strong barrier against mutant invasion.

This analysis suggests the existence of a strong resilience against mutant evolution in the hematopoietic system under biologically realistic parameters, and indicates that more complex conditions need to be present for mutant cell clones to grow. This has implications for the emergence of mutants that give rise to clonal hematopoiesis of indeterminate potential (CHIP), such as *TET2* and *DNMT3A* or *JAK2* mutants. These mutants evolve in the hematopoietic system at homeostasis and increase the chances that hematological malignancies as well as other chronic health conditions develop [20]. For all these mutants, inflammation has been identified as a possible selective force that drives their emergence. In particular, it has been suggested that the mutants themselves induce an inflammatory environment, which negatively affects wild-type cells but to a lesser extent mutant cells, thus creating an increasingly advantageous environment for mutants. We investigated the effect of mutant-induced inflammation on the ability of mutant cells to overcome the invasion barrier in the model. The model suggested that mutant-induced inflammation on its own is unlikely to be sufficient to overcome the invasion barrier, because the mutant cell population first has to increase to a certain extent to induce sufficient inflammation. However, if additionally, aging results in a reduced influx of wild-type stem cells into downstream cell populations through differentiation (as has been shown experimentally [20]), the model suggests that this can reduce the invasion barrier to allow an initially slow mutant growth, which can then accelerate due to rising levels of mutant-induced inflammation. This suggests that complex dynamics are involved in the evolution of mutant cell clones in the hematopoietic system, and provides a guide for experimental testing. A better understanding of these complex dynamics will open doors for the design of evolution-based treatment interventions that can potentially reduce mutant burden and hence alleviate chronic health conditions and reduce the incidence of malignancies.

Our analysis of evolutionary dynamics occurred under the assumption that hematopoietic cell lineages are regulated by feedback control mechanisms during homeostasis, in particular feedback on the self-renewal rate of cells. It is instructive to compare these evolutionary dynamics to those occurring in the absence of feedback control, a setting in which mathematical models parameterized by experimental kinetic data have been analyzed in the past [38]. Without feedback, only carcinogenic mutants can emerge, i.e. cell clones that grow uncontrolled, with a self-renewal rate that is larger than the differentiation rate. Because the self-renewal rate of LT-HSCs is exactly balanced by the differentiation rate, any degree of fitness advantage will result in uncontrolled mutant expansion without negative feedback. Because downstream cell populations (ST-HSCs and MPPs) differentiate more than they self-renew, models without feedback are also characterized by a mutant invasion threshold, although this is simply due to the fact that uncontrolled growth requires the self-renewal rate to exceed the rate of differentiation, which is more difficult to achieve in downstream compartments. The emergence of non-carcinogenic mutants (such as *TET2* or *DNMT3A* mutants in CHIP) is not possible in models without feedback, and the invasion threshold for carcinogenic mutants is then not connected to the influx of wild-type cells from upstream compartments. In contrast, the presence of significant negative feedback in our model ensures that the population remains around a homeostatic steady state as evolution proceeds, and hence non-malignant mutants can emerge, competing with the wild-type cells through the shared feedback inhibition. The mutant invasion barrier for ST-HSCs and MPPs described here arises in our models with feedback due to a different mechanism compared to models without feedback [38]. The invasion threshold in our models with feedback control is connected to competition dynamics, and occurs because the influx of wild-type cells from upstream compartments (and a corresponding lack of influx of mutant cells) confers an inherent disadvantage to the mutant cells, even though they have a higher reproductive fitness in the compartment in which they arise (ST-HSCs or MPPs).

In our model with feedback, the development of uncontrolled cell growth would require further mutations that allow cells to escape feedback-mediated homeostasis [40]. If the degree of negative feedback is significantly weakened (but not absent) in our model, however, the equilibrium level to which invading advantageous mutants grow in the system becomes higher (see Supplement Section 3, Figures S7 and S9), which could represent an intermediate stage in the loss of homeostasis.

While the model structure used in this study is based on the previous literature on modeling the hematopoietic system [21-25] and other tissues [55,56], there are uncertainties regarding the exact processes that should be included in such models for maximal biological realism. All cell divisions in our model are symmetric, and the balance between self-renewal (e.g. a stem cell giving rise to two stem cells) and differentiation (e.g. a stem cell giving rise to two differentiated cells) is determined on the population level. For example, at homeostasis, half of all LT-HSC divisions are self-renewing symmetric divisions, and half are differentiating symmetric divisions. Even though there is empirical evidence for this mechanism in mammalian tissues [57,58], it is possible that asymmetric cell divisions also take place in the hematopoietic system [59], e.g. a stem cell giving rise to one daughter stem cell and one daughter differentiated cell. When asymmetric stem cell divisions are added to the type of models analyzed here, many properties remain the same [60], and we do not expect conclusions to change if asymmetric cell divisions occur among the LT-HSCs. With downstream cells, asymmetric cell divisions are unlikely to play a significant role because at homeostasis, self-renewal and differentiating divisions are not balanced.

A related issue is the mechanism underlying differentiation processes. In our model, differentiation is coupled to cell division. There is experimental evidence that HSC fate decisions are connected to the cell cycle, e.g. through changes in metabolism [61,62], thus supporting our model assumptions. On the other hand, the differentiation of LT-HSCs has been shown to occur in the absence of cell division in certain circumstances [63], although it is not clear how frequently division-independent differentiation events occur. More complex assumptions about differentiation mechanisms can be incorporated into models as more biological information becomes available in the future. Another source of uncertainty concerns the mechanisms that contribute to homeostasis. We modeled feedback in a general way, analyzing a multitude of ways in which feedback control can contribute to homeostasis. Biologically, this can account for both signals that are secreted from the cell lineage itself, and from the microenvironment that senses the abundance of various cell linage sub-populations. Other aspects that might limit the number of cells at homeostasis and that are not part of our model could be spatial constraints [64], as well as interactions with the extracellular matrix [65]. Incorporating these into the current model is a more complex effort and beyond the scope of our study.

The insights obtained from our model can suggest new mouse transplant experiments to test some of these notions, where specific bar code-labeled mutant cells are transplanted together with wild-type cells, and the relative fraction of mutants is monitored over the long-term. Mutant cells can be purified to only include mutant ST-HSCs or MPPs (based on markers), and the fate of the mutants in the different compartment can be tracked. Due to the invasion barriers, we expect the relative fraction of the mutants to decline over time. This experiment could be repeated in the presence of various degrees inflammation, for example induced by LPS. Based on our model, we hypothesize that sufficient inflammation levels help mutants overcome the invasion barrier. This set of experiments could then be repeated, where mutants are present in all cell compartments, including LT-HSCs. Based on our model analysis, we hypothesize that in this case, the mutant fraction will not decline over time even without any inflammation, due to both wild-type and mutant cells being present among the LT-HSCs.

## Notes

### Competing Interest Statement

The authors have declared no competing interest.

