## Supplemental Materials for "Dynamically adjusted cell fate decisions and resilience to mutant invasion during steady state hematopoiesis revealed by an experimentally parameterized mathematical model"

### Supplementary materials

### 1 Wild-type ODE models and their properties

#### 1.1 Model parameterization

We describe hematopoietic turnover by the following system of ODEs:

$$\frac{dx_0}{dt} = r_0 x_0(2p_0 - 1), \quad (1)$$
$$\frac{dx_1}{dt} = 2r_0 x_0(1 - p_0) + r_1 x_1(2p_1 - 1), \quad (2)$$
$$\frac{dx_2}{dt} = 2r_1 x_1(1 - p_1) + r_2 x_2(2p_2 - 1), \quad (3)$$
$$\frac{dx_3^{(1)}}{dt} = 2r_2 x_2(1 - p_2)\gamma^{(1)} - d_3^{(1)} x_3^{(1)}, \quad (4)$$
$$\frac{dx_3^{(2)}}{dt} = 2r_2 x_2(1 - p_2)\gamma^{(2)} - d_3^{(2)} x_3^{(2)}, \quad (5)$$

where the variables $x_0(t), x_1(t)$ and $x_2(t)$ denote the populations of the LT-HSC, SH-HSC, and MPP compartments respectively, and variables $x_3^{(1)}$ and $x_3^{(2)}$ correspond to the CMP and CLP compartments. The corresponding schematic can be found in figure S1(a).

We assume that the probabilities of self-renewal, $p_0, p_1$ and $p_2$ are some functions of the cell populations:

$$p_i = p_i(x_0, x_1, x_2, x_3^{(1)}, x_3^{(2)}), \quad 0 \le i \le 2.$$

We will denote the equilibrium population sizes by capital letters. Solving equations (1-5) in steady state, we obtain:

$$X_1 = \frac{r_0 X_0}{r_1(1 - 2\bar{p}_1)}, \quad (6)$$
$$X_2 = \frac{2r_0 X_0(1 - \bar{p}_1)}{r_2(1 - 2\bar{p}_1)(1 - 2\bar{p}_2)}, \quad (7)$$
$$X_3^{(1)} = \frac{4r_0 X_0(1 - \bar{p}_1)(1 - \bar{p}_2)\gamma^{(1)}}{d_3^{(1)}(1 - 2\bar{p}_1)(1 - 2\bar{p}_2)}, \quad (8)$$
$$X_3^{(2)} = \frac{4r_0 X_0(1 - \bar{p}_1)(1 - \bar{p}_2)\gamma^{(2)}}{d_3^{(2)}(1 - 2\bar{p}_1)(1 - 2\bar{p}_2)}, \quad (9)$$

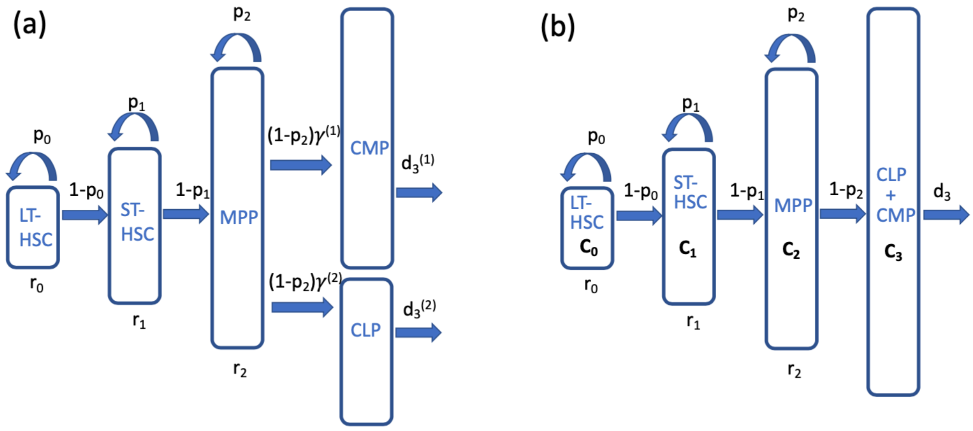

Figure S1: Schematic representations of the compartment system and the processes/rates included in the model. (a) The model with separate CMP and CLP compartments. (b) The model with compartments CMP and CLP combined into compartment $C_3$.

where we denoted

$$\bar{p}_i = p_i(X_0, X_1, X_2, X_3^{(1)}, X_3^{(2)}), \quad 0 \leq i \leq 2,$$

the equilibrium values of the probabilities of self-renewal.

To determine numerical parameters of the model, we used the neutral label propagation data and the relative compartment size data, measured in [1] and [2]. Let us denote the numbers of cells marked with a neutral label as $y_i(t)$ in each of the compartments. The equations for $z_i$ are in this case identical to equations (1-5), and the probabilities of self-renewal /differentiation are evaluated at the equilibrium (and are thus constants).

Denote by $f_i \equiv \frac{z_i}{X_i}$ the fraction of labeled population in each compartment. We have

$$\dot{f}_0 = 0, \quad (10)$$

$$\dot{f}_1 = \frac{1}{\tau_1}(f_0 - f_1), \quad (11)$$

$$\dot{f}_2 = \frac{1}{\tau_2}(f_1 - f_2), \quad (12)$$

$$\dot{f}_3^{(1)} = \frac{1}{\tau_3^{(1)}}(f_2 - f_3^{(1)}), \quad \dot{f}_3^{(2)} = \frac{1}{\tau_3^{(2)}}(f_2 - f_3^{(2)}). \quad (13)$$

where

$$\tau_1 = \frac{1}{r_1(1-2\bar{p}_1)}, \quad \tau_2 = \frac{1}{r_2(1-2\bar{p}_2)}, \quad \tau_3^{(1)} = \frac{1}{d_3^{(1)}}, \quad \tau_3^{(2)} = \frac{1}{d_3^{(2)}}. \qquad (14)$$

The ratios of the compartment sizes at the equilibrium satisfy:

$$\frac{X_1}{X_0} = \frac{r_0}{r_1(1-2\bar{p}_1)}, \quad \frac{X_2}{X_1} = \frac{2r_1(1-\bar{p}_1)}{r_2(1-2\bar{p}_2)}, \quad \frac{X_3^{(1)}}{X_2} = \frac{2r_2\gamma_2^{(1)}}{d_3^{(1)}}, \quad \frac{X_3^{(2)}}{X_2} = \frac{2r_2\gamma_2^{(2)}}{d_3^{(2)}}. \qquad (15)$$

Following the methodology described in [1], we fitted the analytical solutions of equations (10-13) to the time-series of the neutral label proportions in compartments ST-HSC,MPP, CMP, and CLP (using proportions relative to that in the LT-HSC compartment); in addition, the relative compartment sizes were fitted. The prediction for the relative compartment sizes, $\nu_1 = X_1/X_0, \nu_2 = X_2/X_1, \nu_3^{(1)} = X_3^{(1)}/X_2$, and $\nu_3^{(2)} = X_3^{(2)}/X_2$ were obtained from solutions (6-9), and fitted to the quantities reported in [1], which were determined by cell counting of cell suspensions. Note that despite differences in the equations for cell populations used here and in [1], the equations for cell fractions, system (10-13), are identical for our model. In the fitting procedure, we used a larger, updated dataset for the time-series in the SH-HSC and MPP compartments, see [2].

In order to investigate confidence intervals of the fitted parameters, and also the confidence intervals of the fit, we used the bootstrapping method to resample the data, assuming the beta-distribution of the quantities $f_1, f_2, f_3^{(1)}$, and $f_3^{(2)}$ (the usual assumption of normality does not hold in the case where the variables are fractions). The two parameters of the beta-distribution were calculated from the mean and standard deviation information provided in the data. The best fits together with the confidence intervals are shown in figure S2. The results for the best-fitting parameters together with the confidence intervals are presented in Table 1 of the main text.

In what follows we will combine compartments CMP and CLP into a single compartment, which allows for a simpler model, see the schematic of figure S1b. To generalize the description, suppose that there are $n+1$ compartments, $C_0, \ldots, C_n$, that have increasing degree of differentiation (in our case $n = 3$). Denote by $x_i$ the number of wild type cells in compartment

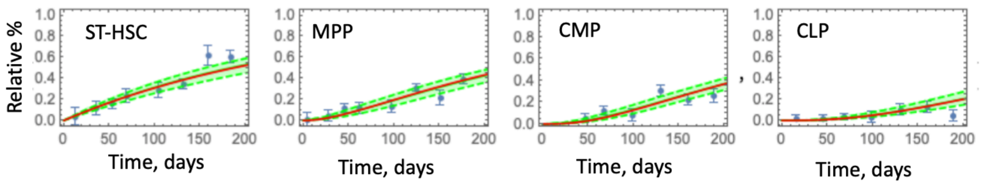

Figure S2: The relative proportions of the neutral label in the four compartments: ST-HSC, MPP, CLP, and CMP. Blue points show the data from [1, 2], with error bars representing the standard error. The red curves are the best fits, and the green shaded area represent the 95% confidence intervals of the fits.

$C_i$. The corresponding system in then given by

$$\frac{dx_0}{dt} = r_0 x_0(2p_0 - 1), \tag{16}$$

$$\frac{dx_i}{dt} = 2r_{i-1}x_{i-1}(1 - p_{i-1}) + r_i x_i(2p_i - 1), \quad 1 \leq i \leq n-1, \tag{17}$$

$$\frac{dx_n}{dt} = 2r_{n-1}x_{n-1}(1 - p_{n-1}) - d_n x_n. \tag{18}$$

Parameter $d_3$ in equation (18) (with $n = 3$) was obtained as $d_3 = (d_3^{(1)}\nu_3^{(1)} + d_3^{(2)}\nu_3^{(2)})/(\nu_3^{(1)} + \nu_3^{(2)})$. The best fit value is $d_3 = 0.0274$ days$^{-1}$, with the 95% C.I. $(0.015, 0.17)$.

#### 1.2 The self-renewal probability functions and model calibration

In system (16-18), the quantity $p_i$ is the probability of self renewal cells in compartment $C_i$. We assume that these are functions of the cell populations. Most generally, we write

$$p_i = p_i(x_0, \ldots, x_n), \quad 0 \leq i \leq n-1. \tag{19}$$

We will assume that these functions are such that a stable, positive steady state

$$x_i(t) = X_n > 0$$

exists. It is given by

$$X_i = X_{i-1}\frac{2r_{i-1}(1-\bar{p}_{i-1})}{r_i(1-2\bar{p}_i)} = X_0\prod_{m=1}^{i}\frac{2r_{m-1}(1-\bar{p}_{m-1})}{r_m(1-2\bar{p}_m)}, \quad 1\le i\le n-1, \quad (20)$$

$$X_n = \frac{2r_{n-1}(1-\bar{p}_{n-1})}{d_n}X_{n-1} \quad (21)$$

(note that $\bar{p}_0 = 1/2$).

The minimal requirements on the functions $p_i$ are as follows:

(i) Functions $p_i$ are probabilities, that is, they satisfy $0 \le p_i \le 1$ for the relevant range of their arguments.

(ii) The self-renewal probabilities are non-increasing functions of the downstream compartment as well as its own compartment, and they are non-decreasing functions of the upstream compartments (whenever applicable).

Here we will consider several special cases of the dependencies of $p_i$ on cell populations. The first three examples below contain two constant nonnegative coefficients per function (those are denoted by $c_i$ and $h_i$). The last example is characterized by a larger number of parameters.

1. *Control from within each compartment,* which we will refer to as "self" for a short-hand notation:

   $$p_i = p_i(x_i), \quad 0\le i\le n-1.$$

   For example, we will use the following functional form:

   $$p_i = \frac{c_i}{1+h_ix_i}. \quad (22)$$

2. *Control from the downstream compartment,* which we will refer to as "next" for a short-hand notation:

   $$p_i = p_i(x_{i+1}), \quad 0\le i\le n-1.$$

   An example is the following functional form:

   $$p_i = \frac{c_i}{1+h_ix_{i+1}}.$$

3. *Control from the most differentiated compartment,* which we will refer to as "last" for a short-hand notation:

$$p_i = p_i(x_n), \quad 0 \le i \le n-1,$$

for example,

$$p_i = \frac{c_i}{1 + h_i x_n}$$

To satisfy constraint (i) above, we will assume that the functional form is relevant in some vicinity of the positive equilibrium, and away from the equilibrium the function is modified such that the probabilities are within $[0, 1]$.

We note that the equilibrium compartment size, equations (20-21), does not depend on our choice of the control strength or type. The constraints on the control parameters come from the self-consistency requirements:

$$p_i(X_0, \dots, X_n) = \bar{p}_i, \quad 0 \le i \le n-1. \tag{23}$$

For the examples of functional forms studied here, if we assume certain values for all the control strength parameters such as $h_i$, equations (23) comprise a linear system system of $n$ equations for $n$ unknowns, $c_0, \dots, c_{n-1}$. For the examples listed above, we have:

1. *Control from within each compartment,* "self":

$$c_0 = \frac{1}{2} + \frac{h_0 \bar{x}_0}{2}, \tag{24}$$

$$c_1 = \bar{p}_1 + \frac{r_0 \bar{x}_0 h_1 \bar{p}_1}{r_1(1 - 2\bar{p}_1)}, \tag{25}$$

$$c_2 = \bar{p}_2 + \frac{2 r_0 \bar{x}_0 h_2 (1 - \bar{p}_1)\bar{p}_2}{r_2(1 - 2\bar{p}_1)(1 - 2\bar{p}_2)}. \tag{26}$$

2. *Control from the downstream compartment,* "next":

$$c_0 = \frac{1}{2} + \frac{h_0 r_0 \bar{x}_0}{2 r_1 (1 - 2\bar{p}_1)}, \tag{27}$$

$$c_1 = \bar{p}_1 + \frac{2 h_1 \bar{p}_1 r_0 \bar{x}_0 (1 - \bar{p}_1)}{r_2(1 - 2\bar{p}_1)(1 - 2\bar{p}_2)}, \tag{28}$$

$$c_2 = \bar{p}_2 + \frac{4 h_2 \bar{p}_2 r_0 \bar{x}_0 (1 - \bar{p}_1)(1 - \bar{p}_2)}{d_3(1 - 2\bar{p}_1)(1 - 2\bar{p}_2)}. \tag{29}$$

3. *Control from the most differentiated compartment,* "last":

$$c_0 = \frac{1}{2} + \frac{2h_0 r_0 \bar{x}_0 (1-\bar{p}_1)(1-\bar{p}_2)}{d_3(1-2\bar{p}_1)(1-2\bar{p}_2)}, \quad (30)$$

$$c_1 = \bar{p}_1 + \frac{4h_1 \bar{p}_1 r_0 \bar{x}_0 (1-\bar{p}_1)(1-\bar{p}_2)}{d_3(1-2\bar{p}_1)(1-2\bar{p}_2)}, \quad (31)$$

$$c_2 = \bar{p}_2 + \frac{4h_2 \bar{p}_2 r_0 \bar{x}_0 (1-\bar{p}_1)(1-\bar{p}_2)}{d_3(1-2\bar{p}_1)(1-2\bar{p}_2)}. \quad (32)$$

Note that these expressions hold in the absence of control, when we simply set $h_i = 0$.

Figure S3 illustrates convergence of the three control models to equilibrium (20-21), by showing the dynamics of the ODEs starting from a small number of cells in the LT-HSC compartment. In all simulations, we fixed control parameters $h_i$ and used the above formulas for the parameters $c_i$. Panel (a) corresponds to the "self" control model, panel (b) to the "next" control model, and panel (c) to the "last" control model. We observe that all three models are capable of reaching the stable equilibrium (values $X_0, \ldots, X_3$, equations (20-21), indicated by dashed horizontal lines in figure S3). These simulations however are not meant to represent the process of development, where different types of feedback control are involved and different parameter values must be used for a realistic description.

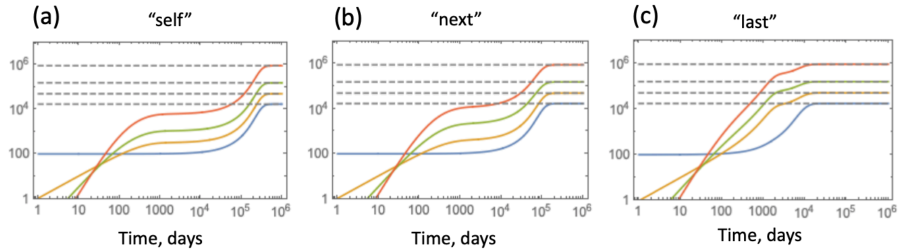

Figure S3: The ODE dynamics of the wild-type system for the three specific assumptions on the control of the self-renewal probabilities: (a) "self", (b)"next", and (c) "last". Populations shown are $x_0$ (blue), $x_1$ (yellow), $x_2$ (green), and $x_3$ (red). The initial conditions are $x_0(0) = 100, x_i(0) = 0$ for $i \geq 1$. The dashed horizontal lines are the equilibrium values, $X_i$, equations (20-21). $h = 10^{-7}$, and the rest of the parameters are as in table 1 of the main text.

### 2 Modeling wild-type and mutant co-dynamics

#### 2.1 ODEs for wild type and mutant cells

In order to introduce mutations in the system, we denote by $y_i$ the mutant population of compartment $C_i$, by $r_i^{(m)}$ the mutant division rates, and by $p_i^{(m)}$ the probabilities of self renewal for mutant cells in compartment $C_i$. Mutants can be generated upon cell division. The four possible division processes for wild type cells, in the presence of a per cell mutation rate $u_i$, are shown in figure S4. Mutants are assumed to divide into mutant daughter cells in the absence of back-mutations or additional mutations. The following set of ODEs describes the co-dynamics of wild type and mutant cells for $n = 3$:

$$\begin{aligned}
\dot{x}_0 &= r_0 x_0 p_0 (1-u_0) - r_0 x_0 (1-p_0), \qquad (33)\\
\dot{y}_0 &= r_0 x_0 p_0 u_0 + r_0^{(m)} y_0 (2p_0^{(m)} - 1),\\
\dot{x}_i &= r_{i-1} x_{i-1} (1-p_{i-1})(2-u_{i-1}) + r_i x_i p_i (1-u_i) - r_i x_i (1-p_i), \quad 1 \le i \le n-1,\\
\dot{y}_i &= r_{i-1} x_{i-1} (1-p_{i-1}) u_{i-1} + 2 r_{i-1}^{(m)} y_{i-1} (1-p_{i-1}^{(m)})\\
&+ r_i x_i p_i u_i + r_i^{(m)} y_i (2p_i^{(m)} - 1), \quad 1 \le i \le n-1,\\
\dot{x}_3 &= r_2 x_2 (1-p_2)(2-u_2) - d_3 x_3,\\
\dot{y}_3 &= r_2 x_2 (1-p_2) u_2 + 2 r_2^{(m)} y_2 (1-p_2^{(m)}) - d_3 y_3. \qquad (34)
\end{aligned}$$

In the absence of mutations ($u_i = 0$), the general terms become:

$$\begin{aligned}
\dot{x}_i &= 2 r_{i-1} x_{i-1} (1-p_{i-1}) + r_i x_i (2p_i - 1),\\
\dot{y}_i &= 2 r_{i-1}^{(m)} y_{i-1} (1-p_{i-1}^{(m)}) + r_i^{(m)} y_i (2p_i^{(m)} - 1).
\end{aligned}$$

In the above model we assume that the differences between wild type and mutant cells could be both in the division rate and in the self-renewal probability. Most generally, we write

$$p_i = p_i(x_0, y_0, \ldots, x_n, y_n), \quad p_i^{(m)} = p_i^{(m)}(x_0, y_0, \ldots, x_n, y_n), \quad 0 \le i \le n-1.$$

A particular formulation that can be used assumes that the mutants' self-renewal probability is a multiple of that of wild-type cells:

$$p_i^{(m)} = (1+s) p_i,$$

where the constant $s$ is the selection coefficient. Alternatively, mutants may be characterized by other differences in their self-renewal function compared to that of wild type cells, such as different control strength coefficients, $h_i$.

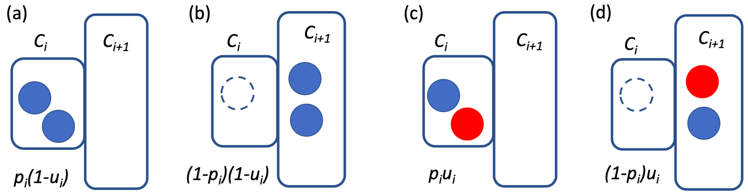

Figure S4: Four types of wild-type cell division: (a) faithful self-renewal, (b) faithful differentiation, (c) self-renewal with a mutation, (d) differentiation with a mutation. The compartments $C_i$ and $C_{i+1}$ are represented by rectangles, wild type cells by blue circles, mutants by red circles, and a position of a cell that has divided and differentiated by dashed circles. The per-cell probabilities of each cell division are indicated for each division type.

#### 2.2 The mutant-free equilibrium and its stability

Consider the main system (33-34), under a general assumption on the control functions:

$$p_i = p_i(x_0 + y_0, x_1 + y_1, x_2 + y_2, x_3 + y_3), \quad 0 \le i \le 2.$$

In the absence of de novo mutations ($u_i = 0, 0 \le i \le 2$), a (positive) mutant-free solution satisfies the algebraic system of equations,

$$\frac{1}{2} = p_0(x_0, \dots, x_3), \tag{35}$$

$$x_1 = \frac{r_0 x_0}{r_1(1 - 2p_1(x_0, \dots, x_3))}, \tag{36}$$

$$x_2 = \frac{2r_0 x_0(1 - p_1(x_0, \dots, x_3))}{r_2(1 - 2p_1(x_0, \dots, x_3))(1 - 2p_2(x_0, \dots, x_3))}, \tag{37}$$

$$x_3 = \frac{4r_0 x_0(1 - p_1(x_0, \dots, x_3))(1 - p_2(x_0, \dots, x_3))}{d_3(1 - 2p_1(x_0, \dots, x_3))(1 - 2p_2(x_0, \dots, x_3))}, \tag{38}$$

$$y_i = 0, \quad 0 \le i \le 3. \tag{39}$$

This system simplifies for our three examples, $p_i = p_i(x_i)$, or $p_i = p_i(x_{i+1})$, or $p_i = p(x_3)$. An explicit solution, if available, will depend on the form of the control functions.

One can investigate stability properties of this mutant-free solution in the general case, by writing out the Jacobian, evaluating it at the equilibrium and determining the eigenvalues, $\lambda_i$, $0 \le i \le 7$. It turns out that there are two distinct groups of eigenvalues. Eigenvalues $\lambda_4, \dots, \lambda_7$ do not depend on the mutant parameters, and the corresponding eigenvectors have zero projections on the 4 mutant directions:

$$\mathbf{v}_i = \begin{pmatrix} v_1^i \\ 0 \\ v_3^i \\ 0 \\ v_5^i \\ 0 \\ v_7^i \\ 0 \end{pmatrix}, \quad 3 \le i \le 7,$$

where the odd components correspond to the $x_0, \dots x_3$ directions and the even components to the $y_0, \dots, y_4$ directions. While the functional shape of the eigenvalues $\lambda_3, \dots, \lambda_7$ depends on the control functions, generally they describe the stability of the positive mutant-free equilibrium in the absence of mutations and can be obtained from the Jacobian of system (16-18). Here we assume that they have a negative real part, that is, a positive equilibrium is stable in the absence if mutants.

The rest of the eigenvalues have a simple form:

$$\lambda_i = r_i^{(m)}(2p_{i,eq}^{(m)} - 1), \quad 0 \le i \le 2, \tag{40}$$

$$\lambda_3 = -d_3, \tag{41}$$

where the functions $p_{i,eq}^{(m)}$ are evaluated at the mutant-free equilibrium. The corresponding eigenvalues have the following form:

$$\mathbf{v}_0 = \begin{pmatrix} v_1^0 \\ v_2^0 \\ v_3^0 \\ v_4^0 \\ v_5^0 \\ v_6^0 \\ v_7^0 \\ v_8^0 \end{pmatrix}, \quad \mathbf{v}_1 = \begin{pmatrix} v_1^1 \\ 0 \\ v_3^1 \\ v_4^1 \\ v_5^1 \\ v_6^1 \\ v_7^1 \\ v_8^1 \end{pmatrix}, \quad \mathbf{v}_2 = \begin{pmatrix} v_1^2 \\ 0 \\ v_3^2 \\ 0 \\ v_5^2 \\ v_6^2 \\ v_7^2 \\ v_8^2 \end{pmatrix}, \quad \mathbf{v}_3 = \begin{pmatrix} v_1^3 \\ 0 \\ v_3^3 \\ 0 \\ v_5^3 \\ 0 \\ v_7^3 \\ v_8^3 \end{pmatrix},$$

From the expressions for the eigenvalues, (40-41), we can see that the mutant division rates (as long as they are positive) do not influence stability. Instead, conditions that guarantee stability of mutant-free solutions involve mutant self-renewal probabilities:

$$p_{i,eq}^{(m)} < 1/2, \quad 0 \le i \le 2. \tag{42}$$

In other words, mutant cells can invade if in at least one of the compartments, they are capable of self-renewal, that is, their self-renewal probability at the wild-type equilibrium exceeds their differentiation probability.

This analysis can be used to examine the fate of mutants originating in different compartments. Still keeping all the mutation rates at zero, assume that mutant production in a given compartment is incorporated through the initial condition:

$$y_i(0) = \begin{cases} \hat{y} > 0, & i = k, \\ 0, & i \neq k, \end{cases} \tag{43}$$

where $k$ is the compartment where the mutation originates. Whether or not this destabilizes the mutant-free solution and leads to a spread of mutants can be determined by examining the relevant eigenvalues.

For mutants originating in $C_0$, all the values in (40) must be negative for stability of the mutant-free state. In other words, as long as any of the three inequalities in (42) are violated, the mutant can invade.

For mutants originating in $C_1$, the initial condition does not have a non-trivial projection onto the eingenvector corresponding to $\lambda_0$ in (40), therefore, only two out of the three conditions in (45) apply, namely, the ones with $i = 1$ and $i = 2$. Consequently, to destabilize the mutant-free solution, one must satisfy the weaker of the two conditions, $p_{i,eq}^{(m)} > 1/2$, $i = 1, 2$.

For mutants originating in $C_2$, only the $i = 2$ condition in (40) is relevant, therefore, to destabilize the mutant-free solution, one must have $p_{2,eq}^{(m)} > 1/2$.

#### 2.3 Fitness thresholds for mutant invasion

Let us consider the special case where the mutant self-renewal probability in each compartment is given by

$$p_i^{(m)} = (1 + s)p_i, \quad 0 \le i \le 2. \tag{44}$$

The quantity $s$ can be positive, zero, or negative, and it can be interpreted as a selection coefficient. Then, stability conditions (42) can be rewritten as

$$(1 + s)\bar{p}_i < 1/2, \quad 0 \le i \le 2. \tag{45}$$

If $\bar{p}_0 = 1/2$ and $\bar{p}_i < 1/2$ for $i = 1$ and $i = 2$, any positive value of $s$ will destabilize the mutant-free solution. Under the assumption in (44), mutant invasion conditions can be formulated as a threshold result. Let us suppose that, in the absence of further de-novo mutations, mutants are placed in compartment $k$, equation (43). Then the mutant invasion conditions become:

$$s > s_c = \min\left\{\frac{1}{2\bar{p}_k} - 1, \ldots, \frac{1}{2\bar{p}_{n-1}} - 1\right\}, \tag{46}$$

that is, the invasion threshold is determined by the equilibrium self-renewal probability in compartment $k$ and all the compartments downstream from it. To summarize: if a mutant is generated in the least differentiated compartment ($C_0$) then any positive value of $s$ is sufficient for the mutant to be advantageous (and invade in the deterministic system). If however the mutant originates in one of the downstream compartments, there is a nontrivial threshold value that $s$ must exceed to be able to invade. For a mutant that originates in compartment $C_k$, the size of the threshold is defined by the largest of the $\bar{p}_l$ values with $l \geq k$, equation (46).

With $n = 3$, there are two cases depending on the relative magnitudes of $\bar{p}_1$ and $\bar{p}_2$. If the equilibrium self-renewal probabilities satisfy

$$1/2 = \bar{p}_0 > \bar{p}_2 > \bar{p}_1, \tag{47}$$

we have the following mutant invasion thresholds:

$$s > s_c = \begin{cases} 0, & k = 0, \\ \min\left\{\frac{1}{2\bar{p}_1} - 1, \frac{1}{2\bar{p}_2} - 1\right\} = \frac{1}{2\bar{p}_2} - 1, & k = 1, \\ \frac{1}{2\bar{p}_2} - 1, & k = 2. \end{cases} \tag{48}$$

This means that for mutants originating in either of the compartments $C_1$ or $C_2$, the same invasion threshold exists, which is the lower of the two values. If, consistent with Table 1 of the main text,

$$1/2 = \bar{p}_0 > \bar{p}_1 > \bar{p}_2, \tag{49}$$

then

$$s > s_c = \begin{cases} 0, & k = 0, \\ \min\left\{\frac{1}{2\bar{p}_1} - 1, \frac{1}{2\bar{p}_2} - 1\right\} = \frac{1}{2\bar{p}_1} - 1, & k = 1, \\ \frac{1}{2\bar{p}_2} - 1, & k = 2. \end{cases} \tag{50}$$

In other words, mutants that originate further downstream will face a higher threshold compared to those that originate lower. It is reasonable to assume that the likelihood of mutant generation in $C_2$ is higher than that for $C_1$, because of the larger size of $C_2$. It therefore appears that the architecture with $\bar{p}_2 < \bar{p}_1$ presents a higher degree of protection against mutant invasion compared to the case with $\bar{p}_1 < \bar{p}_2$.

The value of the self-renewal probability at the equilibrium is responsible for the compartment size: the closer this value is to 1/2, the larger the compartment size (see figure S5(a), where this is illustrated using compartment $C_2$). Therefore, from the point of view of cell amplification in tissue, it is important to keep this quantity closer to 1/2. On the other hand, as this analysis shows, values of $\bar{p}_i$ close to 1/2 tend to lower the mutant invasion threshold for mutants that originate in the given compartment (or even in compartments upstream from that, see figure S5(b)). Therefore, there is a certain trade-off between the functionality and protection against mutations, where the values of self-renewal probability play a pivotal role.

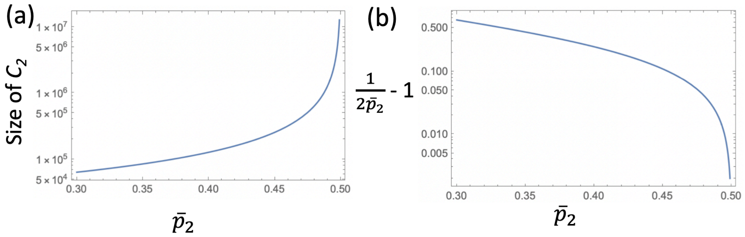

Figure S5: Compartment size (a) and mutant invasion threshold (b) as functions of the equilibrium self-renewal probability $\bar{p}_2$. The rest of the parameters are given in table 1 of the main text.

### 3 The fate of mutants originating in different compartments

In what follows we study how mutants originating in different compartments, may spread through the systems, and how the dynamics depend on the type of control, the strength of the control and and amount of advantage enjoyed by the mutant. We assume that the mutation rate is low, such that once a mutant appears, no new mutations are considered.

##### 3.1 Mutants originating in $C_0$: pure mutant solutions

Since no de-differentiation is assumed in the model, the only way a mutant can occupy compartment $C_0$ is to be generated there. From compartment $C_0$, it can spread to all the downstream compartments. In addition, mutants that originate in downstream compartments, also contribute to the dynamics. If a mutant has an advantage in compartment $C_0$ ($p_{0,eq}^{(m)} > 1/2$) then a purely-mutant solution is established, where the mutants displace the wild-type in $C_0$ and consequently, in all the compartments.

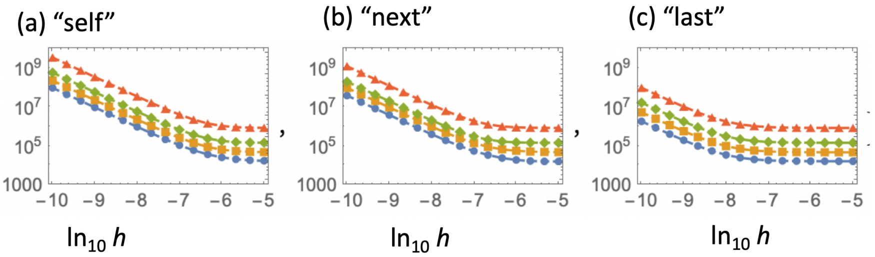

Figure S6: The dependence of the pure mutant equilibrium on the control type and the control strength: (a) "self", (b) "next", and (c) "last" type of control. For each type of control, the equilibrium quantities ($y_0$ (blue), $y_1$ (yellow), $y_2$ (green), $y_3$ (red)) are plotted as functions of $\ln_{10} h$, where $h_i = h$ for all $i$ is control strength. Mutant selection coefficient is $s = 0.01$. The rest of the parameters are given in table 1 of the main text.

While the equilibrium values for the wild-type cells are identical in all of our models of control, mutant behavior depends on both the type of the control functions, and on the strength of control. Figure S6 demonstrates how the pure mutant equilibrium values depend on the strength of control (assuming for simplicity $h_i = h$ for $0 \leq i \leq 2$) for the three types of control system. Generally, the cell numbers for the mutants increase as control decreases, and the solution diverges as $h \to 0$.

Figure S7 shows an example of wild-type (solid) and mutant (dashed) co-dynamics, for different control types (rows) and strengths (columns). The initial condition in each simulation is $y_0(0) = 1$ with other mutant values equal zero at the beginning, and the wild-type values at the mutant-free equilibrium, $x_i(0) = X_i$, $0 \le i \le 3$. As time goes by, we observe that the wild type cells are displaced by mutants. Even though the initial mutant placement is in $C_0$, the compartments turn mutant in the reverse order, from $C_3$ and then $C_2$, followed by $C_1$ and finally $C_0$. This is the consequence of the increasing division rates from least to most differentiated compartments.

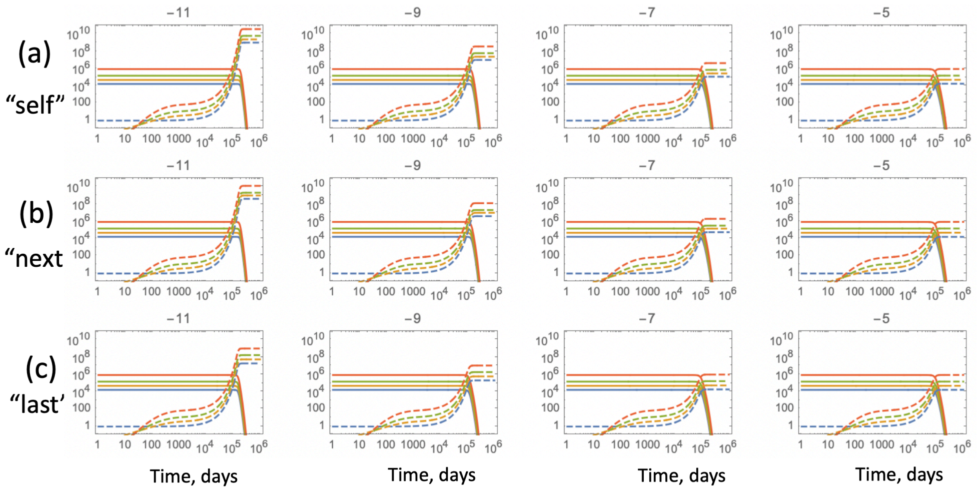

Figure S7: The dynamics of advantageous mutants introduced in $C_0$. The wild type cells (solid lines) and the mutants (dashed lines) in compartments $C_0$ (blue), $C_1$ (yellow), $C_2$ (green), and $C_3$ (red) are plotted as functions of $t$ under the (a) “self”, (b) “next”, and (c) “last” type of control. The different panels in each row correspond to different control strengths, with the quantity $\ln_{10} h$ marked in curly brackets above each plot. $s = 0.01$, $u_i = 0$ for all $i$, and the rest of the parameters are given in table 1 of the main text.

#### 3.2 Mutants originating in downstream compartments

As was mentioned previously, if a mutant originates in compartment $C_0$, under any positive value of $s$ it will act as an advantageous mutant. The situation is different if the mutant originates in one of the downstream compartments. In order to destabilize the mutant-free solution, the mutant must be able to self-renew (at the mutant-free equilibrium conditions) in the compartment of its origin or at least one of the downstream compartments. Under model (44) this translates to a threshold condition for the selection coefficient

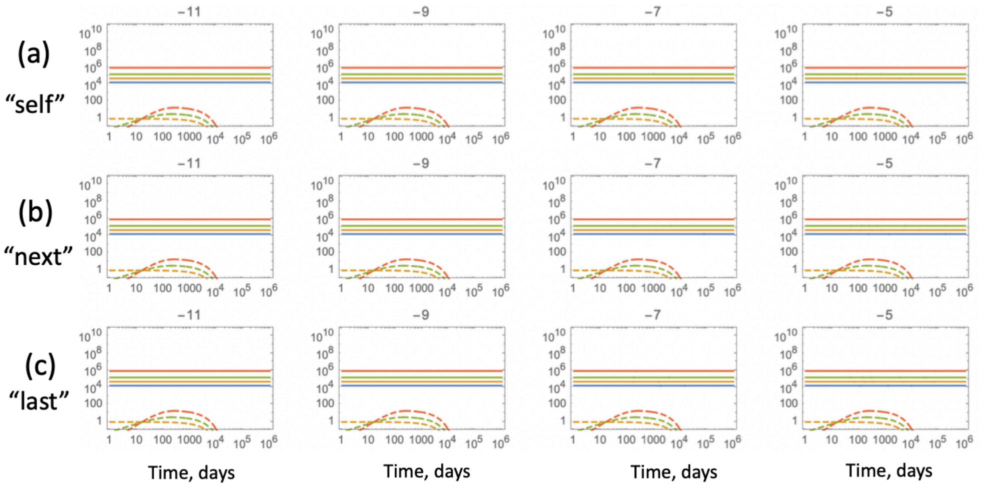

Figure S8: The dynamics of advantageous mutants ($s = 0.05$) introduced in $C_1$. Mutant fitness is below the threshold: $s < s_c^{(1)} \approx 0.058$. The notations and the rest of the parameters are as in figure S7.

$s$. For the parameters of table 1 of the main text, we have

$$s_c^{(1)} \equiv \frac{1}{2\bar{p}_1} - 1 \approx 0.058, \quad s_c^{(2)} \equiv \frac{1}{2\bar{p}_2} - 1 \approx 0.203.$$

Since the self-renewal rates satisfy inequality (49), the invasion thresholds are given by (50). Simulations below demonstrate this result.

In figures S8 and S9, the mutant introduced in $C_1$ and has selection coefficient $s = 0.05$ and $s = 0.06$, respectively. In figure S8, we have $s < s_c^{(1)}$, that is, the selection coefficient is below the threshold in $C_1$. As a result, the mutant dies out in its compartment of origin ($C_1$, yellow dashed lines) and is unable to spread. In contrast to this, in figure S9 we have $s_c^{(2)} > s > s_c^{(1)}$, that is, the selection coefficient exceeds the threshold in $C_1$, and a mutant population gets established in $C_1$. Even though the threshold for compartment $C_2$ is not reached by these mutants, the existence of the input from $C_1$ makes the threshold in $C_2$ irrelevant, and the mutants subsequently spread to the downstream compartments.

As with mutants originating in $C_0$, the steady state level of successfully spreading mutants depends on the strength of the feedback, measured by $h$ here. For high $h$ (strong feedback) the mutants' equilibrium is lower than

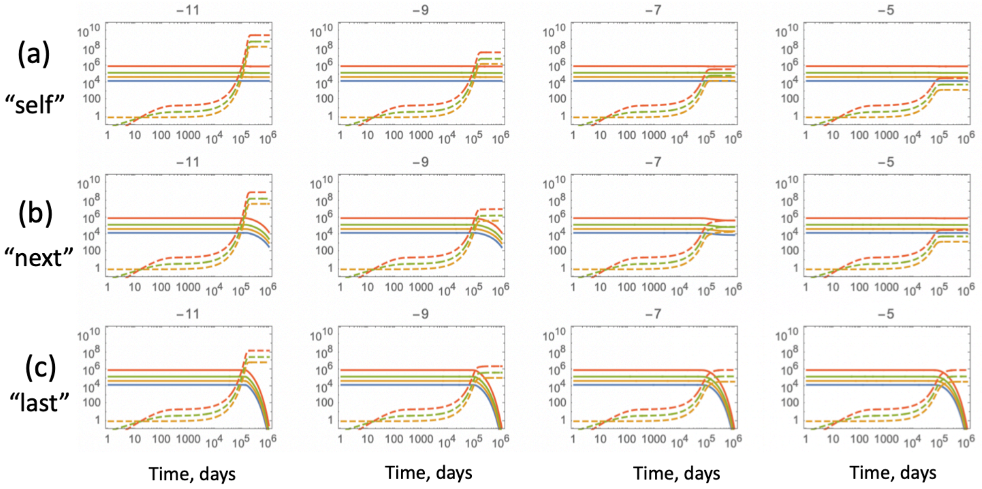

Figure S9: The dynamics of advantageous mutants ($s = 0.06$) introduced in $C_1$. Mutant fitness is above the threshold: $s > s_c^{(1)} \approx 0.058$. The notations and the rest of the parameters are as in figure S7.

the wild-type equilibrium in the respective compartments (see the rightmost panels in figure S9). For weaker values of feedback, the equilibrium value of the mutants grows (and diverges for $h \to 0$).

Figure S9 demonstrates another difference compared to mutants originating in $C_0$: in cases where the mutant is introduced in $C_1$, the resident wild-type population is not necessarily driven extinct by the expanding mutant, resulting in coexistence steady states. The fate of wild-type cells depends on the model formulation for the control of self-renewal probabilities. In the model where the control is "self" type (panel (a)), the wild type populations remain constant and the mutants grow to their equilibrium values. If the feedback is of type "next" or "last" (panels (b) and (c)), the wild-type populations may become decreased or go extinct in the presence of mutants.

The timing of mutant invasion depends on the amount of mutant advantage. Figure S10 compares the dynamics of mutants with $s = 0.2$ (significantly over the invasion threshold in $C_1$) with those with $s = 0.06$ (just above the threshold, depicted in figure S10). Apart from an enormous acceleration in the mutant rise in all models, we also observe a higher level of the mutant equilibrium.

If mutants originate in $C_2$, under the parameter values of Table 1 of the

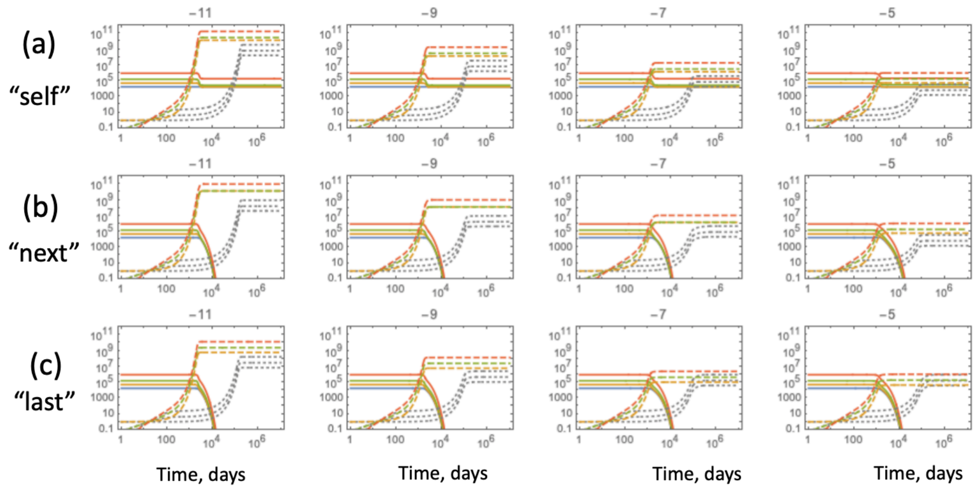

Figure S10: The dynamics of advantageous mutants with a larger fitness ($s = 0.2$, compared to figure S9) introduced in $C_1$. The mutant numbers under the assumption of $s = 0.06$ (same as those in figure S9) are shown in dotted gray lines, for comparison. The notations and the rest of the parameters are as in figure S7.

main text, they have a much higher threshold to overcome. If the value $s < s_c^{(2)}$, the mutants will die out. Figure S11 shows mutants with $s = 0.21$, just above the threshold. These mutants successfully spread. The patterns of dependence on the feedback stricture and strength are similar to what was noted above.

#### 3.3 Control of "self" type, mutants originating in $C_1$: a case study

Here we examine a specific system with mutants originating in $C_1$, and focus our attention on the equilibria in that compartment. In the absence of mutations, we have for $C_1$:

$$\dot{x}_1 = r_0 x_0 + r_1 x_1 (2p_1 - 1), \quad (51)$$

$$\dot{y}_1 = r_1 y_1 (2(1+s)p_1 - 1); \quad (52)$$

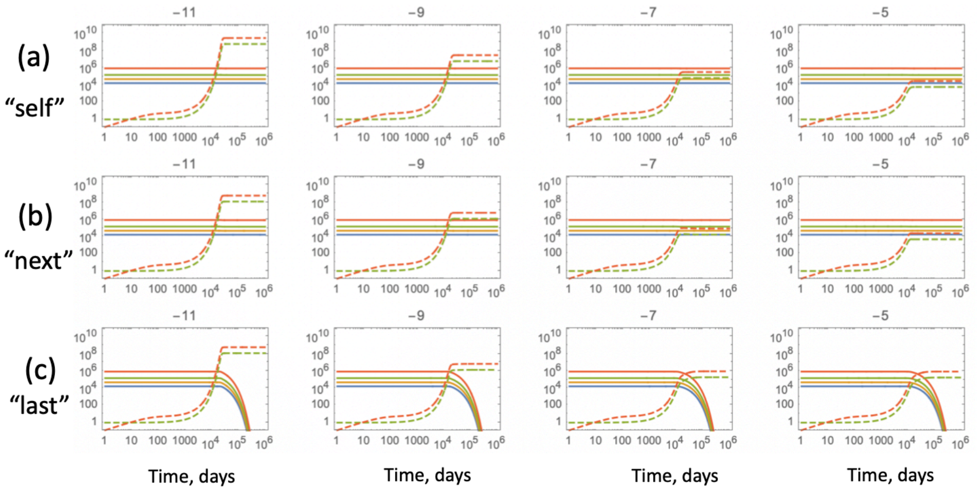

Figure S11: The dynamics of advantageous mutants ($s = 0.21$) introduced in $C_2$. Mutant fitness is above the threshold: $s > s_c^{(2)} \approx 0.203$. The notations and the rest of the parameters are as in figure S7.

note that we took $p_0(x_0) = 1/2$, which remains constant in this model, where $p_i = p_i(x_i + y_i)$. It is useful to write down the Jacobian of the system:

$$J = \begin{pmatrix} r_1(2p_1 - 1) + 2r_1x_1p_1' & 2r_1x_1p_1' \\ 2r_1y_1(1+s)p_1' & r_1(2(1+s)p_1 - 1) + 2r_1y_1(1+s)p_1' \end{pmatrix}. \quad (53)$$

If $x_0p_0 > 0$, there are two steady state solutions, discussed below.

**The mutant-free solution:** $y_1 = 0$, and $x_1$ is given by

$$x_1 = \frac{r_0x_0}{r_1(1 - 2\bar{p}_1)}. \quad (54)$$

The eigenvalues of the corresponding Jacobian are $r_1(2p_1 - 1) + 2rx_1p_1'$ and $r_1(2(1+s)p_1 - 1)$, evaluated at the steady state. This solution is stable when $(1+s)p_1 < 1/2$, and becomes unstable if $(1+s)\bar{p}_1 > 1/2$. As the value of $s$ grows, the system experiences a transcritical bifurcation at

$$s = s_c = \frac{1}{2\bar{p}_1} - 1. \quad (55)$$

As explained for the general case, for $0 < s < s_c$, mutants, although having a positive selection coefficient, effectively behave as disadvantageous mutants. The threshold value is higher if $\bar{p}_i$ is further from $1/2$.

**Mutant solution:** $y_1 > 0, x_1 > 0$, $(1+s)p_1(x_1, y_1) = 1/2$, and

$$x_1 = \frac{r_0 x_0 (1+s)}{r_1 s}. \tag{56}$$

The number of wild type cells is independent of control, but the number of mutants depends strongly on control, increasing as the control parameter becomes weaker. This solution becomes stable when $s$ is greater than critical ($s > s_c$). This solution is characterized by mutants that are able to maintain their numbers in the compartment, through a sufficiently high value of $s$. For the particular shape of control defined in (22), the steady state is given by equation (56) and

$$y_1 = \frac{(2\bar{p}_1(1+s) - 1)(r_1 s(2\bar{p}_1 - 1) - h_1 r_0 x_0 (1+s))}{h_1 r_1 s (2\bar{p}_1 - 1)}. \tag{57}$$

While $x_1$ is independent of control, $y_1$ depends strongly on $h_1$. There are two distinct regimes, separated by the value $h_c$,

$$h_c = \frac{s r_1 (1 - 2\bar{p}_1)}{r_0 x_0 (1+s)}.$$

- *Weak control.* If $h_1 \ll h_c$, we have

  $$y_1 \approx \frac{2\bar{p}_1(1+s) - 1}{h_1}, \tag{58}$$

  that is, it is inversely proportional to control parameter $h_1$, and tends to infinity in the absence of control ($h_1 \to 0$).

- *Strong control.* If $h_1 \gg h_c$, we have a constant ($h_1$-independent) level of mutants,

  $$y_1 \approx \frac{r_0 x_0 (1+s)(2\bar{p}_1(1+s) - 1)}{r_1 s (2\bar{p}_1 - 1)}. \tag{59}$$

Figure S12 shows the behavior of solution (57) and its approximations.

### 4 Including replication limits in the lineage dynamics

#### 4.1 The wild type system

Let us suppose that LT-SCs can divide indefinitely, but cells in the downstream compartments have a replication limit, which we call $K$. Accordingly,

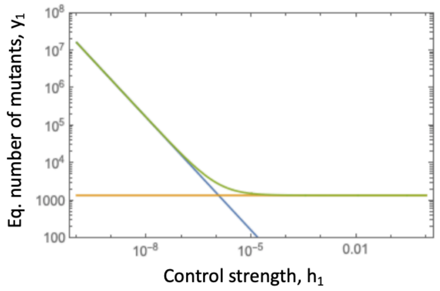

Figure S12: The equilibrium number of mutants (originating in $C_1$), equation (57), green, as a function of the control strength, $h_1$. Blue and orange lines represent approximations (58) and (59), respectively. Here $s = 0.06$, and the rest of the parameters are given in table 1 of the main text.

for compartments $C_i$ with $i = 1$ and $i = 2$, we will split all cells in division classes, such that $x_1 = \sum_{k=1}^{K} x_{1k}$, $x_2 = \sum_{k=1}^{K} x_{2k}$; here the second index, $k$, enumerates the division classes, while the first index stands for the compartment number. We will assume that cells differentiate from compartment $C_0$ into compartment $C_1$ by entering class $x_{11}$. For divisions in class $x_{ik}$, at each cell division, the progeny is placed in class $x_{i,k+1}$ with probability $p_i$ (a self-renewal division), and in class $x_{i+1,k+1}$ with probability $1 - p_i$ (a differentiation division). We assume that the progeny of cells in classes $x_{iK}$ is removed from the system. The following of ODEs describes these processes:

$C_0$ :

$$\dot{x}_0 = r_0 x_0 (2p_0 - 1), \tag{60}$$

$C_1$ :

$$\begin{aligned} \dot{x}_{11} &= 2r_0 x_0 (1 - p_0) - r_1 x_{11}, \\ \dot{x}_{1k} &= 2r_1 x_{1,k-1} p_1 - r_1 x_{1k}, \quad 2 \le k \le K, \end{aligned}$$

$C_2$ :

$$\begin{aligned} \dot{x}_{21} &= -r_2 x_{21}, \\ \dot{x}_{22} &= 2r_1 x_{11}(1 - p_1) - r_2 x_{22}, \\ \dot{x}_{2k} &= 2r_1 x_{1,k-1}(1 - p_1) + 2r_2 x_{2,k-1} p_2 - r_2 x_{2k}, \quad 3 \le k \le K. \end{aligned}$$

$C_3$ :

$$\dot{x}_3 = 2r_2 \sum_{k=1}^{K} x_{2k}(1 - p_2) - d_3 x_3. \tag{61}$$

We will assume, as before, that the probabilities of self-renewal are functions of the quantities $x_i$, $i \in \{0,1,2,3\}$. At the equilibrium, we have $p_i = \bar{p}_i$, with $\bar{p}_0 = 1/2$. The equilibrium solution is then given by:

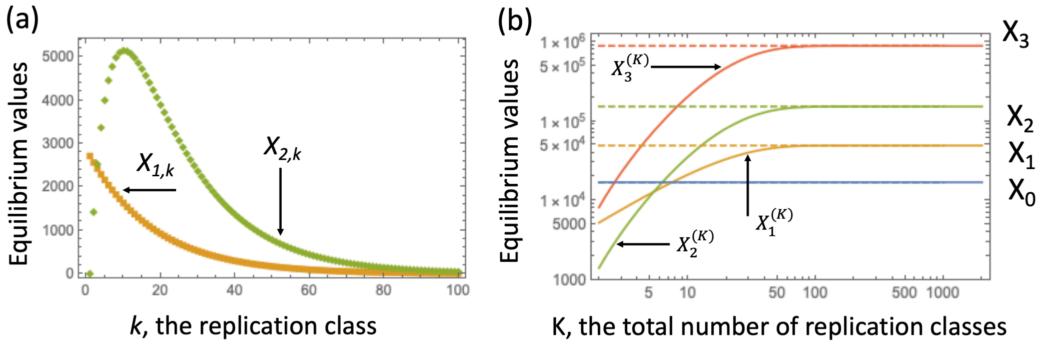

Figure S13: Equilibrium solutions for the system with replication limits. (a) Equilibrium values in the individual replication classes, $X_{1,k}$ and $X_{2,k}$, equations (63-64). (b) The equilibrium values of the cell populations in the 4 compartments as functions of the total number of replication classes, $K$. The horizontal dashed lines show the equilibrium values for the basic system in the absence of replication limits, equations (20-21).

$$X_0 = \bar{x}_0, \tag{62}$$

$$X_{1,k} = \frac{r_0 \bar{x}_0}{r_1}(2\bar{p}_1)^{k-1}, \quad 1 \le k \le K, \tag{63}$$

$$X_{2,1} = 0,$$

$$X_{2,k} = \frac{r_1}{r_2} X_{11} 2^{k-1}(1-\bar{p}_1)\sum_{m=0}^{k-2} \bar{p}_1^m \bar{p}_2^{k-2-m} = \frac{r_0 \bar{x}_0}{r_2} 2^{k-1}(1-\bar{p}_1)\frac{\bar{p}_1^{k-1} - \bar{p}_2^{k-1}}{\bar{p}_1 - \bar{p}_2}, \tag{64}$$

$$X_3^{(K)} = \frac{2r_2(1-\bar{p}_2)}{d_3}\sum_{k=1}^{K} X_{2,k} =$$

$$\frac{2r_0\bar{x}_0(1-\bar{p}_1)(1-\bar{p}_2)}{d_3}\,\frac{2(\bar{p}_1-\bar{p}_2) + (2\bar{p}_2)^K(1-2\bar{p}_1) - (2\bar{p}_1)^K(1-2\bar{p}_2)}{(\bar{p}_1-\bar{p}_2)(1-2\bar{p}_1)(1-2\bar{p}_2)}. \tag{65}$$

For the total populations of the two middle compartments, we have

$$X_1^{(K)} = \sum_{k=1}^{K} X_{1,k} = \frac{r_0\bar{x}_0}{r_1}\frac{1-(2\bar{p}_1)^K}{1-2\bar{p}_1}, \quad (66)$$

$$X_2^{(K)} = \sum_{k=1}^{K} X_{2,k} = \frac{r_0\bar{x}_0(1-\bar{p}_1)}{r_2}\,\frac{2(\bar{p}_1-\bar{p}_2)+(2\bar{p}_2)^K(1-2\bar{p}_1)-(2\bar{p}_1)^K(1-2\bar{p}_2)}{(\bar{p}_1-\bar{p}_2)(1-2\bar{p}_1)(1-2\bar{p}_2)}. \quad (67)$$

Since both $2\bar{p}_1 < 1$ and $2\bar{p}_2 < 1$, we have

$$\lim_{K\to\infty} X_1^{(K)} = X_1, \quad \lim_{K\to\infty} X_2^{(K)} = X_2, \quad \lim_{K\to\infty} X_3^{(K)} = X_3,$$

see equations (20-21). Figure S13(a) shows the equilibrium values in different replication classes, $X_{1,k}$ and $X_{2,k}$. The total equilibrium populations of the four compartments (equations (66-67)) are shown in figure S13(b), as functions of the total number of replication classes, $K$. We observe that they converge to the values obtained previously for the system in the absence of replication classes. The convergence for $X_1^{(K)}$ is faster than that for $X_2^{(K)}$, as the latter is defined by $2\bar{p}_2 > 2\bar{p}_1$. The equilibrium values of the compartment sizes increase with $K$, the total number of replication classes, see figure S13(b). If the number of replication classes is greater than about 100, the system can achieve amplification for the cell numbers of increasing degrees of differentiation.

To ensure the match of system (60-61) under a specific choice of feedback on self-renewal probability, $p_i(x_0,\dots,x_3)$, we use equations (23), where $X_1 = \sum_{k=1}^{K} X_{1,k}$ and $X_2 = \sum_{k=1}^{K} X_{2,k}$. For the numerical examples below we used the specific form of control, $p_i = c_i/(1+h_i(x_i+y_i))$ (the "self" type). In this case, equations (23) yield:

$$c_i = \bar{p}_i(1+h_iX_i), \quad 0 \le i \le n-1.$$

#### 4.2 Co-dynamics of wild-type and mutant cells.

In order to include mutations, denote by $y_0$, $y_{1,k}$, $y_{2,3}$ and $y_3$ (with $k$ enumerating the replication classes as before) the populations of the different types of mutants. We can write a cascade of equations for the mutants, which is similar to that of the wild-types (equations (60-61)), coupled through both the process of mutations and the self-renewal probabilities, which are functions of both wild-type and mutant populations. If a mutation occurs in compartment $C_0$, we observe patterns similar to those described previously: any advantageous mutant will displace wild-type cells in compartment $C_0$ and consequently, in the whole lineage. It is more interesting to consider mutant generation in the downstream compartments.

For this purpose, it is convenient to ignore de-novo mutations. The equations for the wild-type cells remain the same as (60-61), except now $p_i = p_i(x_0, y_0, \ldots, x_3, y_3)$. The equations for the mutant cells look similar, except they contain information about the difference between mutant and wild-type kinetics. We will consider several types of mutations.

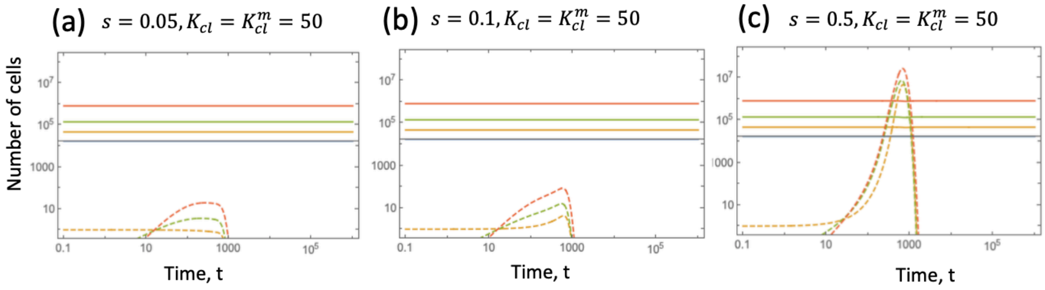

Figure S14: Mutant and wild-type dynamics for the system with replication limits, where mutations do not affect replication limits:the role of fitness advantage. The initial conditions are given by equations (62-65) for the wild-types, and all the mutant classes are zero except $y_{1,1}(0) = 1$. Total populations $x_i$ and $y_i$, $0 \leq i \leq 3$, are plotted as functions of time. Wile-type (mutant) numbers are shown by solid (dashed) lines in compartment $C_0$ (blue), $C_1$ (yellow), $C_2$ (green), and $C_3$ (red). (a) $s = 0.05$, (b) $s = 0.1$, (c) $s = 0.5$. Under the "self" feedback model, $h = 10^{-9}$, $u = 0$, $K = 50$, and the rest of the parameters are as in table 1 of the main text.

**Mutations that do not affect the replication capacity.** Let us assume that as before, mutations affect only the self-renewal probabilities of the cells, which is reflected in the factor $1 + s$. From studying the system in the absence of replication limits, we learned that mutants cannot rise if their fitness advantage, $s$, is below a threshold, specific to the compartment of origin. We also know that once $s$ is above the threshold, the rise of the mutant population occurs faster for larger values of $s$.

These patterns remain in the system with replication limits, but there is in additional factor that plays a role in mutant dynamics. Mutants that originate in any upstream compartment, have a finite life-span during which they can rise, and once their replication capacity is exhausted, they are removed from the system. Therefore, any rise/domination of mutants can only occur for a limited amount of time. There is no steady state where mutants prevail, in the absence of mutant cells residing in $C_0$.

As a result, the mutants' ability to make an impact on the system depends on their ability to rise to significant numbers, before they are wiped out due to reaching the replication capacity. Figure S14 demonstrates this by plotting the numbers of both wild-type (solid lines) and mutant (dashed lines) cells in a system with replication capacity $K = 50$, where initially, a single mutant cells is introduced in compartment $C_1$ (type $y_{1,1}$). All the parameters are the same for the three panels except for the mutant fitness parameter, $s$. In panel (a), the mutant fitness advantage $s$ is below the invasion threshold ($s < s_c^{(1)} \approx 0.058$), resulting in the number of mutants in $C_1$ declining. Note that before all mutants die out, they give rise to a certain number of cells in the downstream compartments, but being below the threshold prevents the mutant from expansion. In panel (b), the mutant fitness is above the threshold, but the time before the replication capacity of mutants is exhausted is too short for the mutants to make a significant impact. In panel (c), the mutant fitness is significantly higher and they rise much faster, such that they reach large numbers before they are wiped out.

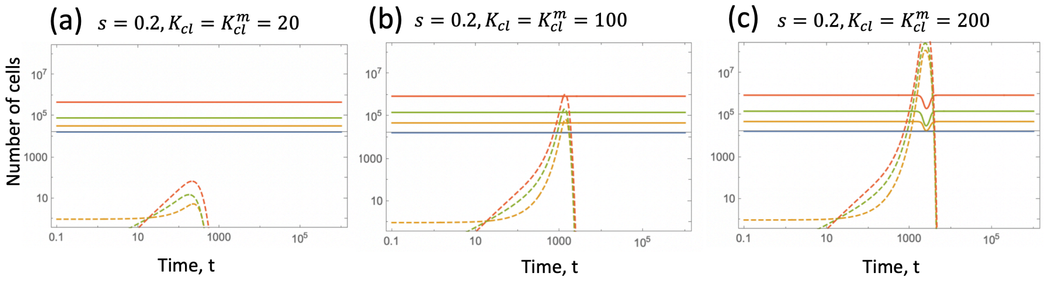

Figure S15: Mutant and wild-type dynamics for the system with replication limits, where mutations do not affect replication limits: the role of the replication limits. (a) $K = 20$, (b) $K = 100$, (c) $K = 200$. In all panels, $s = 0.2$. Notations and the rest of the parameters are as in figure S14.

In contrast to figure S14, in figure S15 the mutant fitness parameter, $s$, is kept constant, but the replication capacity of all cells is varied. The larger the replicative capacity, the longer is the time period during which the mutants can expand. In all panels of figure S15, we have $s > s_c^{(1)}$, and the replication capacity takes values $K = 20$, $K = 100$, and $K = 200$ in panels (a)-(c), respectively. The mutant's impact is the highest in panel (c), where it has a chance to rise to significant levels before crashing to zero.

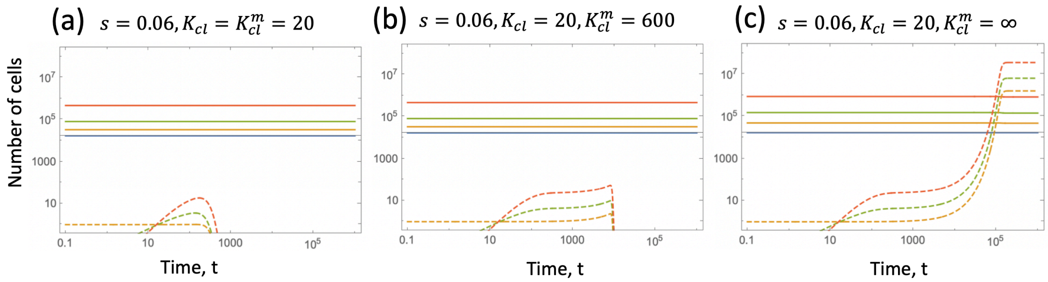

Figure S16: Mutant and wild-type dynamics for the system, where mutations affect both self-renewal probability and the replication limits. (a) $K_m = 20$, (b) $K_m = 600$, (c) $K_m = \infty$. In all panels, $s = 0.06$ and $K = 20$. Notations and the rest of the parameters are as in figure S14.

**Mutations that do affect the replication capacity.** If the mutations are assumed to increase the replication capacity of the cells, the time during which they can expand increases, and therefore the maximum level achieved by the mutants also increases. In figure S16, the carrying capacity of the wild-type cells is kept at $K = 20$, and mutants are just above the invasion threshold ($s = 0.06$). In panel (a), the mutants do not have an increased replication limit, and disappear before having a chance to expand. In panel (b), the replication capacity of mutants is increased with respect to that of the wild-types, resulting in a longer mutant presence. In panel (c) we assume that mutants can divide indefinitely, and observe the establishment of a mutant steady state.

The same trends are observed if the mutant is introduced in compartment $C_2$ instead of $C_1$. In figure S17, a mutant generated in $C_2$ with the fitness advantage just above the threshold ($s = 0.21 > s_c^{(2)} \approx 0.203$) does not have a chance to rise if the replication capacity is low (panel (a)), but increasing the mutants' carrying capacity ($K_m = \infty$ in panel (b)) leads to the establishment of a mutant equilibrium. In panel (c), the mutant's fitness advantage, $s$, is higher, such that it rises to significant levels even under a finite replicative capacity ($K_m = 200$), before being flushed of the system.

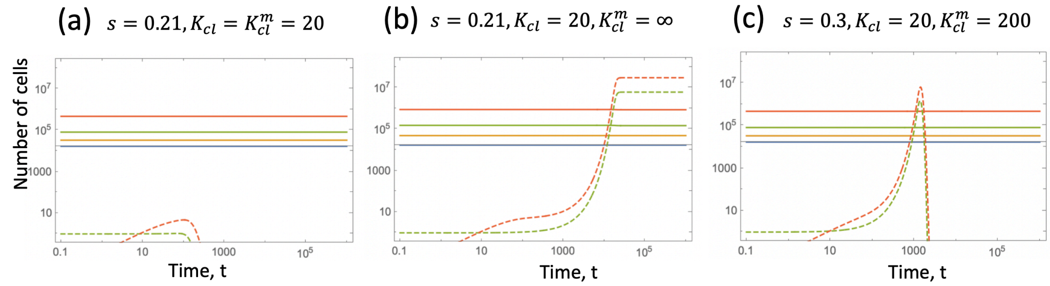

Figure S17: Similar to figures S14,S15, and S16, but the mutant originates in compartment $C_2$. Initially, all the mutant classes are zero except $y_{2,2}(0) = 1$. (a) $s = 0.21, K = K_m = 20$, (b) $s = 0.21, K = 20, K_m = \infty$, (c) $s = 0.3, K = 20, K_m = 200$. Notations and the rest of the parameters are as in figure S14.
